# Gender imbalance in citations most pronounced in high-impact neuroscience journals

**DOI:** 10.1101/2025.02.19.638905

**Authors:** Kat Hefter, Erin G. Teich, Dani S. Bassett

## Abstract

In the past several years, neuroscience, like many other fields, has worked to address pervasive gender imbalances. Although tangible improvements have been made in academic publishing and conference participation, gender imbalances in citations among leading neuroscience journals persist and are increasing with time. Here, we expand upon prior work by providing a more comprehensive analysis of citation practices across fifty journals to improve our understanding of the dynamics of the field as a whole, particularly across different kinds of journals. We first confirm that reference lists tend to include more papers with men as first and last author than expected if gender were unrelated to citation practices. Further, we demonstrate that this effect is strongest in the most influential journals. We also find that the apparent closure of the gender gap is due to the interaction of three unique, nuanced patterns of citation: one that is characterized by relative overcitation of papers with men as both first and last author; one that is characterized by relative overcitation of women as the last author, and one that is characterized by relative undercitation of women as the first author. In addition, we find that these patterns of citation are not distributed evenly across the field: more prestigious journals tend to have articles with citation gaps favoring men, while less prestigious journals tend to overcite papers with women as last authors. We discuss potential drivers for these patterns and suggest some implications for the field moving forward.

## Introduction

Gender bias is a well-documented phenomenon in the scientific community. Inequalities exist in a range of contexts, including but not limited to compensation (1), hiring and promotions (2–4), authorship (5), teaching evaluations (6, 7), grant funding (8, 9), collaboration credit (10), publication quantity (11), journal tier (12), and number of citations (13, 14).

Given the importance of academic visibility on career advancements (15, 16), attempts to mitigate these disparities often take the form of detecting and addressing gender imbalances in avenues to recognition (17). Indeed, due in large part to the work of groups such as BiasWatchNeuro (http://biaswatchneuro.com/), Women in Neuroscience (http://winrepo.org/), and Anne’s List (http://anneslist.net/), among others, there is some evidence that the gender gap in speaking opportunities is closing (18). In addition, the field may be seeing some improvement in publishing opportunities (13), and some journals are taking steps to have more diverse and representative communities of authors and reviewers (19).

Despite these improvements, however, gaps still exist in the number of citations accrued by scholars. Reference lists of publications tend to include more papers with men as first and last authors than expected given the gender composition of the field, a pattern shown not only in neuroscience (13, 14), but also physics (20), astronomy (21), international relations (22, 23), political science (24), and medicine (25), to name a few. Given the importance of citation metrics to an author’s perceived prestige, and the potential negative down-stream effects of inequitable engagement, understanding the extent and drivers of these imbalances is crucial.

One key component of this effort is to understand how universal gender imbalances are. Because prior work has focused primarily on the most prestigious journals (e.g., (13, 25)), the relationship between gender imbalance and journal prestige has not been well-studied. In addition, at least one study has found that citation bias may be mediated by author gender distribution (26). Given gender-based disparities in tier of journal publication (12), the interaction between journal prestige, author gender distribution, and citation practices merits further exploration.

Therefore, in this study, we take a more holistic look at the field of neuroscience by quantifying gender imbalance in citation in articles published in the top fifty neuroscience journals as defined by the Eigenfactor score. Within the pool of papers published between 1995 and 2023, we build a network of citing-cited papers, and locate and remove self-citations. We then study gender-dependent citation behavior, according to the following questions. First, we ask whether women-led papers are cited at the same rates as we would expect, given their other relevant characteristics like author seniority or time since publication. Next, we test whether there are any differences in undercitation between men-led and women-led reference lists. While we initially sought to simply determine whether results from the most prestigious journals extended to the field at large, we find instead that citation behavior is more complex: specifically, we identify three subgroups of journals, with differing author composition and journal prominence, that display opposing patterns of citation behavior. One subgroup shows a pattern identical to that of the more prominent journals in our previous study: overcitation of papers with men as both first and last author. A second subgroup tends to show a relative overcitation of women as the last author, and the third is characterized by relative undercitation of women as the first author. In addition, these groups are not distributed equally across the field: the overcitation of men occurs more in more prestigious journals, while the overcitation of women as last author tends to occur more in in less prestigious journals. Our results motivate further efforts to excavate and understand differential drivers of citation imbalances.

## Results

### Data collection and author gender modeling

We extracted data from the Thomson Reuters’ Web of Science (WoS) database for research articles and reviews published in the 50 top neuroscience journals since 1995, based on their Eigenfactor scores (27) (see Supplementary Table 1 for the full list of journals). Analyses were conducted using an open-source codebase (13) that has been used in several studies with similar methodologies (14, 20, 26, 28–30). In all, 362,772 articles were included in the dataset of cited and citing papers.

We assigned each author a probabilistic gender based on their first name, using the methods described in Ref. (13) (see Methods for more details). The label ‘man’ was assigned to authors whose first name had a ≥ 0.7 chance of belonging to someone identifying as a man, and the label ‘woman’ was assigned to authors whose first name had a ≥ 0.7 chance of belonging to someone identifying as a woman. We performed our analyses using the articles for which gender could be assigned with high probability for both the first and last author (78%, *n* = 281, 776).^1^ Assigning gender labels in this way created four categories for our analyses: man as first and last author [MM]; woman first author, man last author [WM]; man first author, woman last author [MW]; and woman first and last author [WW].

We note that our ability to assign gender labels based on first names is inherently limited. Besides the probabilistic uncertainty and the fact that many names can be used by people of all genders, authors may be intersex, transgender, or nonbinary. Our model fails to account for these other gender possibilities. However, our model usefully mimics the experience of human citers, who often lack personal knowledge of the gender of the authors they cite. Hence, our model enables us to study bias not only based on the known gender of the author, but also on the inferred gender of the author.

### Calculation of citation imbalance

When quantifying citation behavior within neuroscience articles, we only considered papers published in any of these top 50 journals (Supplementary Table 1) between 2009 and 2023 (*n* = 230, 798 in total, *n* = 183, 713 for articles for which gender could be assigned with high probability for both the first and last author). The 2009 starting year was chosen both to allow for comparison of our results to prior work (13), and to increase the chances that the citing paper’s cited references fell within our broader dataset. Thus, while any paper in the dataset could be a *cited* paper, only papers published after 2009 were analyzed as *citing* papers.

For each citing paper, we took the subset of its citations that had been published in one of the top 50 journals since 1995, removed self-citations (defined as any cited papers where either the first or last author of the citing paper was first or last author of the cited paper), and calculated the probability that each of the cited papers fell into the four author categories described above. Single-author papers by men and women were included in the MM and WW categories, respectively. Of the 2,011,704 citations made between 2009 and 2023, 1,167,080 (58.0%) were made to MM papers, 472,277 (23.5%) were made to WM papers, 203,765 (10.1%) were made to MW papers, and 168,582 (8.4%) were made to WW papers.

We calculated citation imbalance using two metrics: (1) the Literature-Based Citation Gap and (2) the Conditioned Citation Gap. Schematics of both methods can be found in Fig. 1. Each citation gap is calculated by subtracting the percent expected from the percentage observed, and then dividing this difference by the percentage expected. The two approaches differ in the way that they calculate the expectation.

**Figure 1.**
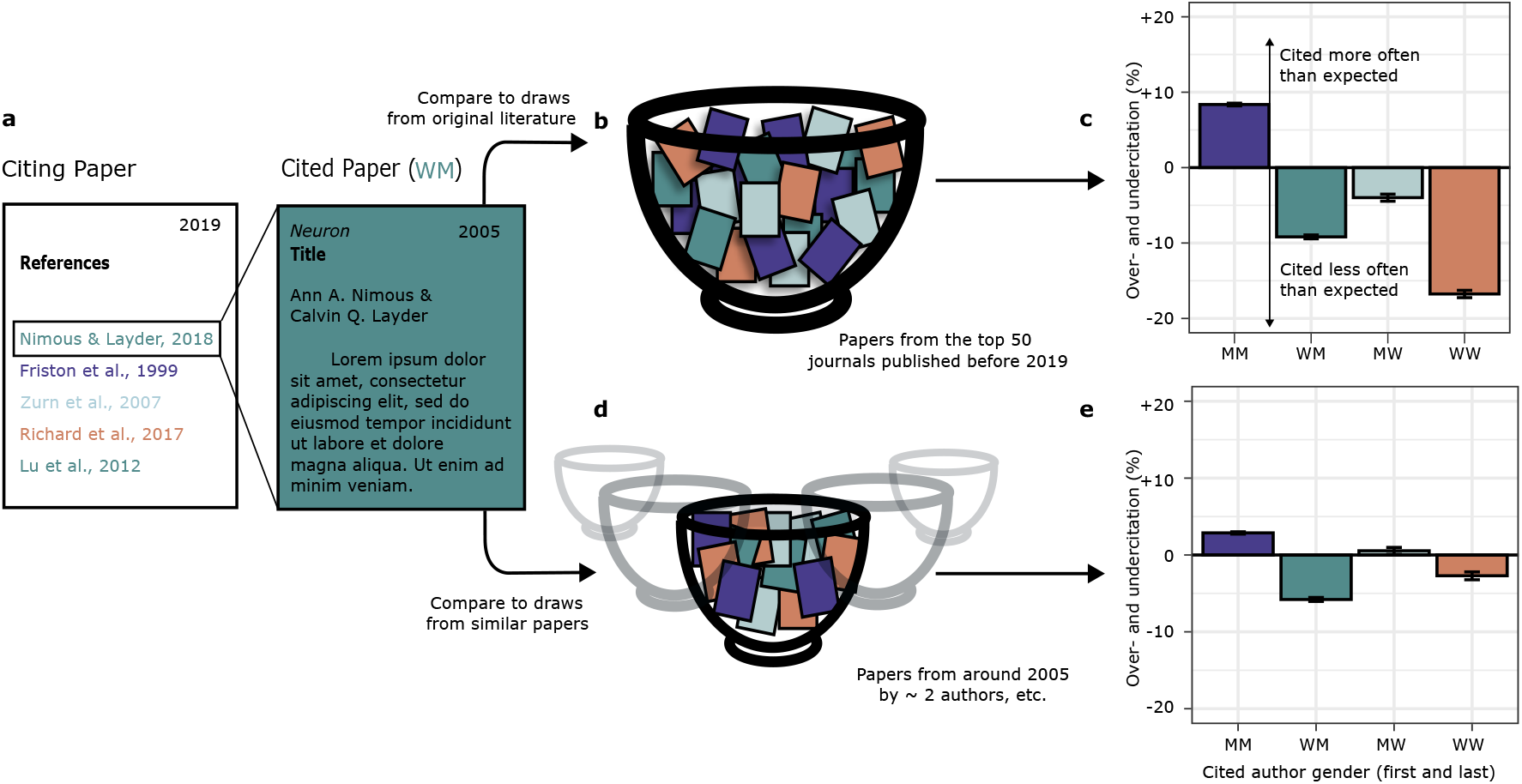
Schematic for comparison of different citation gap models. (a) All of the papers cited in a given paper’s reference list are classified into one of four gender categories. **(b-c) Literature-based model:** (b) The proportion of papers in each gender category are compared to the gender proportions of existing literature at the time of publication. (c) The gap between proportions of gender groups observed compared to expectation. **(d-e) Conditional model:** (d) gender proportions in reference lists are compared to gender proportions of the subset of articles published that match relevant characteristics across multiple domains. (e) The difference between the proportion of gender groups observed versus expected. Bars represent overall over- and undercitation according to a given model, calculated from 2,011,704 total citations. Error bars represent the 95% CI of each over- and undercitation estimate, calculated from 500 bootstrap resampling iterations.

The **Literature-Based Citation Gap** compares the observed number of citations in each gender category to the number that would be expected if references were drawn randomly from the pool of papers that existed prior to the citing paper (Fig. 1a.). For each paper, we calculated the gender proportions among all papers published prior to the citing paper, then multiplied by the number of papers cited. We then summed these values across all papers. The expected proportions based solely on the pool of citable papers were 53.5% for MM, 25.9% for WM, 10.5% for MW, and 10.1% for WW. Using this measure, MM papers were cited 8.4% more than expected (95% CI=(8.2%, 8.6%), one-sample *t* = 89.34, df = 499, *p* < 0.0001), WM papers were cited 9.2% less than expected (95% CI=(−9.5%, -8.9%), one-sample *t* = 67.96, df = 499, *p* < 0.0001), MW papers were cited 4.0% less than expected (95% CI=(−4.5%, -3.5%), one-sample *t* = 18.62, df = 499, *p* < 0.0001), and WW papers were cited 16.8% less than expected (95% CI=(−17.2%, -16.2%), one-sample *t* = 65.19, df = 499, *p* < 0.0001).

The Literature-Based Citation Gap method has proven useful for deriving insight but also somewhat limited in that it does not account for other relevant properties of published papers that might affect the rate at which they are cited (i.e., when and where the paper was published, number of authors and author seniority, and whether the article was a research article or a review). We therefore also modeled the **Conditioned Citation Gap** with the same formula as above, but using a generalized additive model (GAM) to calculate expected proportions based on these potentially relevant factors. Specifically, we used the (1) month and year of publication; (2) combined number of publications by first and last author; (3) number of total authors on the paper; (4) journal of publication; and (5) whether the paper was a review (see Methods, detailed descriptions in Ref. (13), and Fig. 1b.). Once again, the expected gender proportions were calculated for each citing paper and then multiplied by the number of papers cited. These were then summed to create the total expected proportions. For the remainder of this paper, we focus on the conditional citation gap, and for brevity refer to it simply as the *citation gap*.

Based on the relevant properties of cited papers, the new expected citation rates were 56.4% for MM, 25.0% for WM, 10.1% for MW, and 8.6% for WW. Thus, after accounting for salient non-gender characteristics, MM papers were still cited 2.8% more than expected (95% CI=(2.7%, 3.0%), one-sample *t* = 29.59, df = 499, *p*<0.0001), WM papers were cited 5.8% less than expected (95% CI=(−6.1%, -5.5%), one-sample *t* = 33.85, df = 499, *p* < 0.0001), and WW papers were cited 2.7% less than expected (95% CI=(−3.2%, -2.1%), one-sample *t* = 8.11, df = 499, *p* < 0.0001). Of 2,011,704 total citations, these values correspond to citations being given to MM papers around 33,500 more times than expected, compared to approximately 27,400 fewer times for WM papers and 4,600 fewer times for WW papers. MW papers did not show a significant citation gap (one-sample *t* = 1.02, df = 499).

Interestingly, these statistically significant gender gaps were smaller in magnitude than those previously reported for the top five journals alone; Dworkin *et al*. (2020) reported conditional gaps of 5.2%, -6.7%, -4.6%, and -13.9% for MM, WM, MW, and WW teams, respectively (13). We therefore next sought to determine whether the difference in magnitude was due to our expanded sample size, and more precisely, whether citation gaps tracked with journal prominence.

### The effect of journal prominence on citation behavior

Previous work on this topic only explored the trends in authorship in the top five neuroscience journals at the time by Eigenfactor score (*Neuron, Nature Neuroscience, Neuroimage, Journal of Neuroscience*, and *Brain*). To determine whether those trends were representative of the field as a whole, we analyzed both the publishing and citation rates of both the top five journals and the remaining forty-five journals. We found that the top five journals were highly prolific: despite making up only a tenth of the journals studied, they accounted for roughly one-fifth of all papers published between 2009 and 2023. Further, the proportion of MM-authored papers was higher, and decreased more slowly, in the more prestigious journals than the field as a whole. In 1995, 66.8% of the papers published in the top five journals were by MM authors, compared to 65.3% of the field as a whole; in 2023, 42.5% of the papers published that year in the top five journals were by MM authors, compared to 39.0% for the field as a whole (Fig. 3). In addition, the average yearly increase in percentage of papers published by non-MM teams in the top five journals was only 0.70% each year (95% confidence interval (CI) = 0.63%, 0.77%)), compared to 0.98% each year (95% confidence interval (CI) = 0.94%, 1.03%)) for the field as a whole.

The top five journals also played a significant role in accounting for field-wide citation imbalances. Specifically, within reference lists from the top 5 journals (*n* = 40, 425 papers), MM papers were cited 7.1% more than expected (95% CI = (6.8 %, 7.4%), one-sample *t* = 44.44, df = 499, *p* < 0.0001), WM papers were cited 10.8% less than expected (95% CI = (−11.4%, -10.3%), one-sample *t* = 38.46, df = 499, *p* < 0.0001), MW papers were cited 4.6% less than expected (95% CI = (−5.5 %, -3.6%), one-sample *t*-statistic = 9.39, df = 499, *p* < 0.0001), and WW papers were cited 15.0% less than expected (95% CI = (−16.2%, -13.9%), one-sample *t*-statistic = 25.94, df = 499, *p* < 0.0001). These gaps matched or exceeded previous findings (13).^2^ In contrast, within the reference lists of papers not within the top five journals (*n* = 143, 288 papers), the citation gap nearly disappeared or even switched directions. MM papers were cited only 1.1% more than expected (95% CI = (0.9%, 1.3%), one-sample *t* = 10.49, df = 499, *p* < 0.0001), WM papers were cited 3.9% less than expected (95% CI = (−4.3%, -3.6%), one-sample *t* = 21.92, df = 499, *p* < 0.0001), MW papers were cited 2.2% *more* than expected (95% CI = (1.5 %,2.8 %), one-sample *t* = 6.82, df = 499, *p* < 0.0001), and WW papers were cited 1.7% *more* than expected (95% CI = (1.0%, 2.3%), one-sample *t* = 4.67, df = 499, *p* < 0.0001)).

The observed differences between reference lists from the top five journals and reference lists from the remaining forty-five journals, calculated using the bootstrapped estimates of the gap values, were all significant (Welch two-sample *t*-test: MM: *t* = 696.53, df = 872.15, *p* < m.p., WM: *t* = -463.49, df = 847.99, *p* < m.p., MW: *t* = -258.5, df = 864.77, *p* < m.p., WW: *t* = -548.12, df = 835.12, *p* < m.p.). These results are summarized in Fig. 2(a-c).

**Figure 2.**
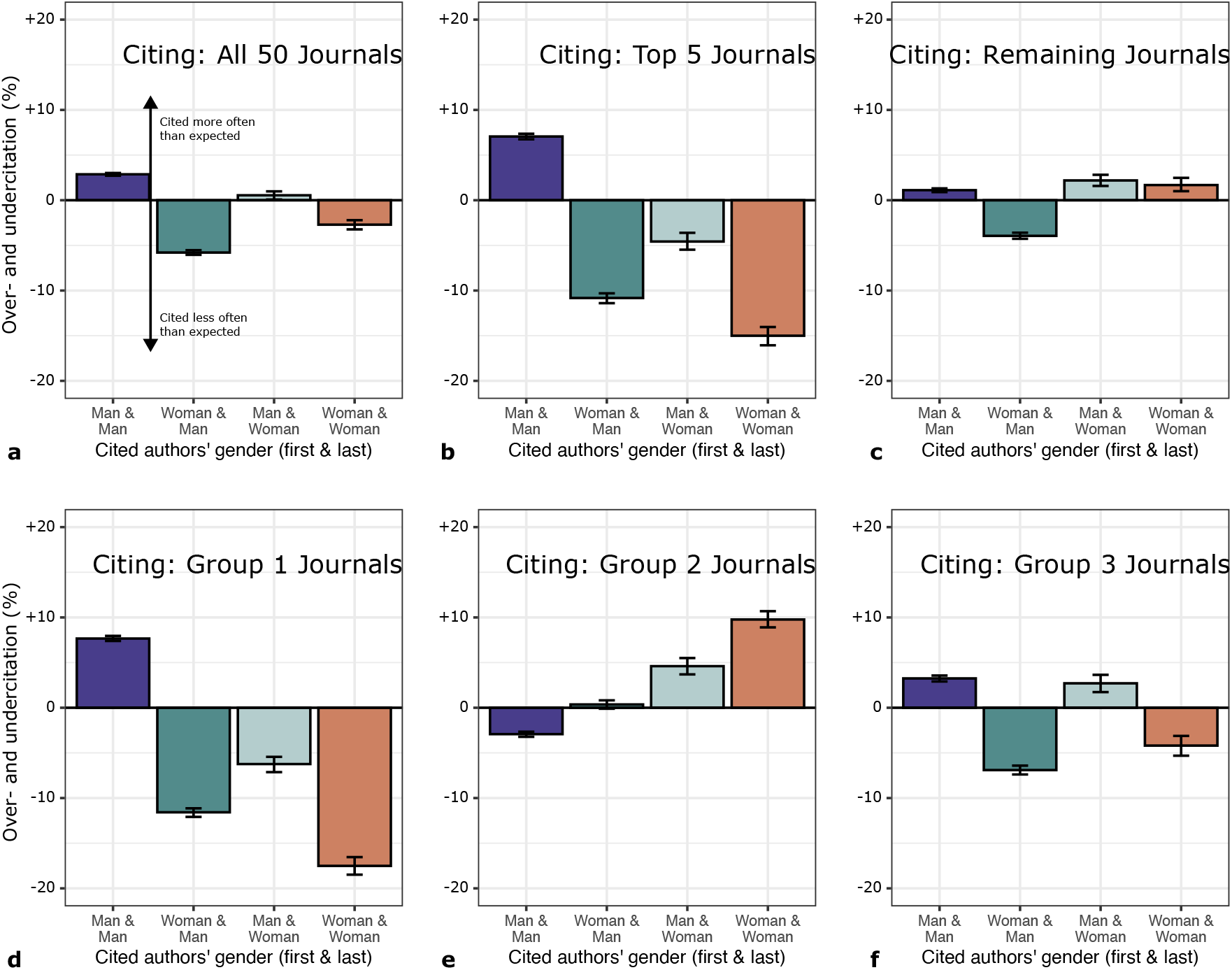
Different sets of journals showed different patterns of citation gaps. (a) Citation gap for the field as a whole, versus (b) citation gap for the top 5 journals and (c) citation gap for the remaining 45 journals. (d-f) Citation gaps for the journals in each of the three clusters identified. Group 1 is characterized by overcitation of MM articles and undercitation of all other categories (and closely resembles the citation gap for the top 5 journals); Group 2 is characterized by overcitation of papers with a woman as the last author, and Group 3 generally shows undercitation of papers with a woman as first author.

**Figure 3.**
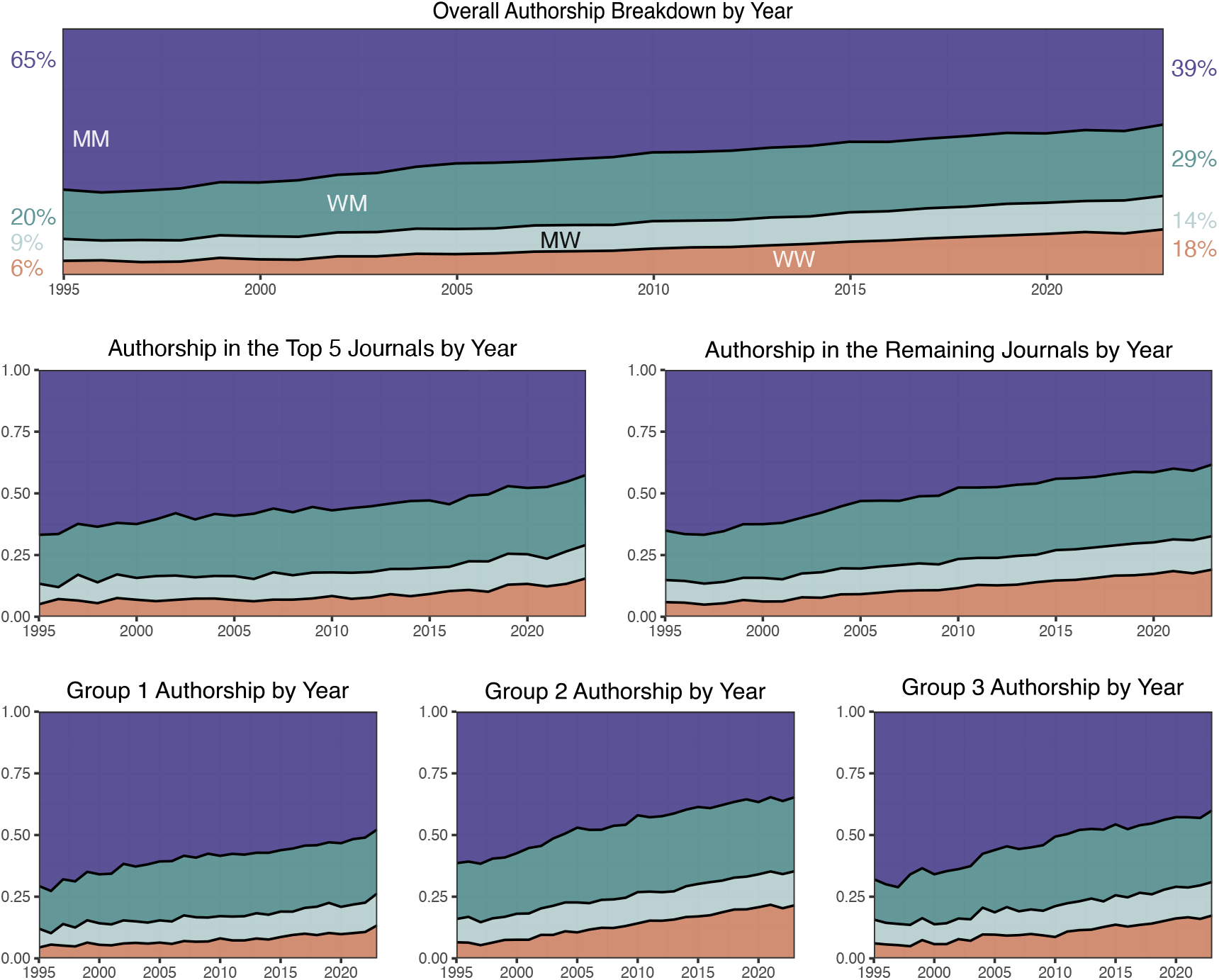
Trends in gender composition of authorship remain generally consistent across subgroups. Overall trends across all fifty journals studied (top), the top five journals versus the remaining forty-five journals (middle), and the three subgroups of journals (bottom). Graphs show the relative proportions of articles with men as the first and last author (MM, purple), women as first author and men as last author (WM, green), men as first author and women as last author (MW, grey), and women as first and last author (WW, orange).

### Differential patterns of citation activity

To explore whether there was something unique about the top journals, we sought to determine whether journals could be separated into categories with different citation activity. First, we calculated the citation gap separately for papers originating from each journal. Then, we used a *K*-means clustering method to group the journals based on their citation gaps. This approach yielded three clusters of 12, 22, and 16 journals, respectively. The number of clusters was chosen heuristically to maximize cluster stability and minimize the number of singleton clusters (see Supplementary Information for details). Journals in each of these clusters showed markedly different features. Group 1 contained the most prestigious journals, had a higher percentage of men authors, and was more likely to overcite men and undercite women. Group 2 contained more of the less-prestigious journals, with a higher proportion of women authors, and overcited women last authors. Finally, Group 3 was the fastest-growing group, with a faster-growing proportion of women-authors and a slight gender gap that disfavored women first authors. In addition, a qualitative inspection of the titles of papers in each of the three groups seemed to suggest some topic-based stratification (See Extended Data Figures 10,11, and 12) for more details.) The citation gaps for each of these groups can be seen in Fig. 2(d-f).

### Group 1: MM Overcitation

This group, which contained 12 journals and accounted for 28.4% of the citing articles, was characterized by overcitation of MM articles and undercitation of articles in every other category. In this cluster, MM papers were cited 7.6% more than expected (95% CI = (7.4%, 8.0%), one-sample *t* = 54.12, df = 499, *p* < 0.0001), WM papers were cited 11.6% less than expected (95% CI = (−12.1%, -11.1%), one-sample *t* = 47.39, df = 499, *p* < 0.0001), MW papers were cited 6.2% less than expected (95% CI = (−7.1 %, -5.4%), one-sample *t* = 13.43, df = 499, *p* < 0.0001), and WW papers were cited 17.5% less than expected (95% CI = (−18.5%, -16.5%), one-sample *t* = 34.55, df = 499, *p* < 0.0001) (Fig. 2(d)). Notably, four of the top five journals (*Neuron, Nature Neuroscience, Neuroimage*, and *Journal of Neuroscience*), and five of the top ten journals, were members of this category.

### Group 2: _W Overcitation

This group was the most populous group, containing 22 journals and 43.7% of the citing articles. It was characterized by overcitation of papers with a woman as the last author, and a slight undercitation of MM papers. In this cluster, MM papers were cited 2.9% *less* than expected (95% CI = (−3.2%, -2.7%), one-sample *t* = 20.18, df = 499, *p* < 0.0001), MW papers were cited 4.6% more than expected (95% CI = (3.7%, 5.5%), one-sample *t* = 10.48, df = 499, *p* < 0.0001), and WW papers were cited 9.8% more than expected (95% CI = (8.9%, 10.7%), one-sample *t* = 21.01, df = 499, *p* < 0.0001). There was not a significant gap for WM papers (one-sample *t* = 1.52, df = 499) (Fig. 2(e)). Only one of the journals in this category was in the top 10, at number 10.

### Group 3: W_ Undercitation

The final group, which contained 16 journals and made up 27.9% of the total pool of papers, was characterized by undercitation of papers with first authors who were women, and a slight overcitation of papers with first authors who were men. In this cluster, MM papers were cited 3.2% more than expected (95% CI = (2.9%, 3.6%), one-sample *t* = 20.11, df = 499, *p* < 0.0001), WM papers were cited 6.9% less than expected (95% CI = (−7.4%, -6.4%), one-sample *t* = 26.81, df = 499, *p* < 0.0001), MW papers were cited 2.7% more than expected (95% CI = (1.7 %, 3.6%), one-sample *t* = 5.29, df = 499, *p* < 0.0001), and WW papers were cited 4.2% less than expected (95% CI = (−5.3%, -3.1%), one-sample *t* = 7.33, df = 499, *p* < 0.0001) (Fig. 2(f)). Four of these journals were in the top ten journals in the field.

For each gender category, a one-way ANOVA showed a significant effect of group membership on the gender citation gap (df = 2, *F* = {634931, MM; 298911, WM; 75756, MW; 349458, WW}). Tukey’s HSD post hoc test showed that MM groups were cited most in Group 1, then Group 3, then Group 2 (*p* < m.p. in all cases), and WM, MW, and WW groups were all cited most in Group 2, then Group 3, then Group 1 (*p* < m.p. in all cases). In addition, there was a relationship between group membership and prestige, as measured through the Eigenfactor score (27). A one-way ANOVA revealed a significant effect of group membership on journal eigenfactor score (df = 2, *F* = 4.442, *p* = 0.0171), and a post-hoc Tukey PSD found that Group 1 (MM overcitation) was significantly more prestigious than Group 2 (_W overcitation) (p=0.012). Group 3 (W_ undercitation) fell between the other two and was not significantly different in prestige from either (*p* = 0.16, Group 3 < Group 1, and *p* = 0.53, Group 3 > Group 2).

### Citation groups differ by gender composition

Groups notably differed in their publishing trends. The majority of articles published in Group 1 journals were written by MM teams (55%); in addition, the proportion of articles written by MM teams published each year remained higher than both other groups from 1995 to 2023. Articles in Group 2 journals were more likely to be written by teams with at least one woman as first or last author (61%), and consistently had lower proportions of MM articles published each year than the other two groups. Group 3 journals began with similar publishing proportions as Group 1, but had a faster growth in the proportion of articles published with a woman as first or last author; approximately 55% of the pool of citing papers from Group 3 had at least one woman.

The proportion of articles published in each group has also changed over time, as summarized in Fig. 3. The proportion of articles published in Group 3 journals has risen steadily from around 14% in 1995 to 38% in 2023. By contrast, the proportion of articles published in Group 1 journals has steadily decreased from around 50% in 1998 to 17% in 2023. Group 2 articles have hovered around 45% of the total articles.

### The relation between authors’ gender and citation behavior

Previous research in the fields of neuroscience (13), physics (20), political and social science (24), international relations (22), and communication (29), among others, has shown that undercitation of women-led papers occurs to a greater extent within men-led reference lists. To test whether this held true for the neuroscience field as a whole, and for any of its subgroups, we compared the gender makeup of references within papers that had men as both first and last author (MM) to those within papers that had at least one woman as an author (W∪W, including WM, MW, and WW papers).

### Entirety of the field

We first examined whether there were consistent patterns of citation as a function of gender category, irrespective of citation group. Of the 183,713 articles published in the top 50 journals between 2009 and 2023, 44.7% were MM and 55.3% were W ∪ W. As we observe in Figure 4, papers led by men first and last authors largely accounted for the overcitation of man-led papers and undercitation of women-led papers. Specifically, within MM reference lists (*n* = 82, 169 papers), other MM papers were cited 6.9% more than expected (95% CI = (6.7%, 7.1%), one-sample *t* = 55.25, df = 499, *p* < 0.0001), WM papers were cited 9.7% less than expected (95% CI = (−10.1%, -9.3%), one-sample *t* = 46.66, df = 499, *p* < 0.0001), MW papers were cited 4.0% less than expected (95% CI = (−4.8 %, -3.3%), one-sample *t* = 10.26, df = 499, *p* < 0.0001), and WW papers were cited 14.9% less than expected (95% CI = (−15.7%, -14.1%), one-sample *t* = 35.58, df = 499, *p* < 0.0001). Within W∪W reference lists (*n* = 101, 544 papers), MM papers were cited 0.5% less than expected (95% CI = (−0.7 %, -0.3%), one-sample *t* = 4.15, df = 499, *p* < 0.0001), WM papers were cited 2.9 % less than expected (95% CI = (−3.2 %, -2.5%), one-sample *t* = 14.72, df = 499, *p* < 0.0001), MW papers were cited 3.8% more than expected (95% CI = (3.1 %, 4.5%), one-sample *t* = 10.42, df = 499, *p* < 0.0001), and WW papers were cited 6.8% more than expected (95% CI = (6.0%, 7.6%), one-sample *t* = 16.76, df = 499, *p* < 0.0001). Welch two-sample t-tests comparing citation gaps for each gender category across MM- and W ∪ W-led teams were all significant (MM: one-sample *t* = -964.69, df = 994.04, *p* < m.p.; WM: one-sample *t* = 536.87, df = 993.59, *p* < m.p.; MW: one-sample *t* = 326.33, df = 990.65, *p* < m.p.; WW: one-sample *t* = 833.66, df = 996.73, *p* < m.p.).

**Figure 4.**
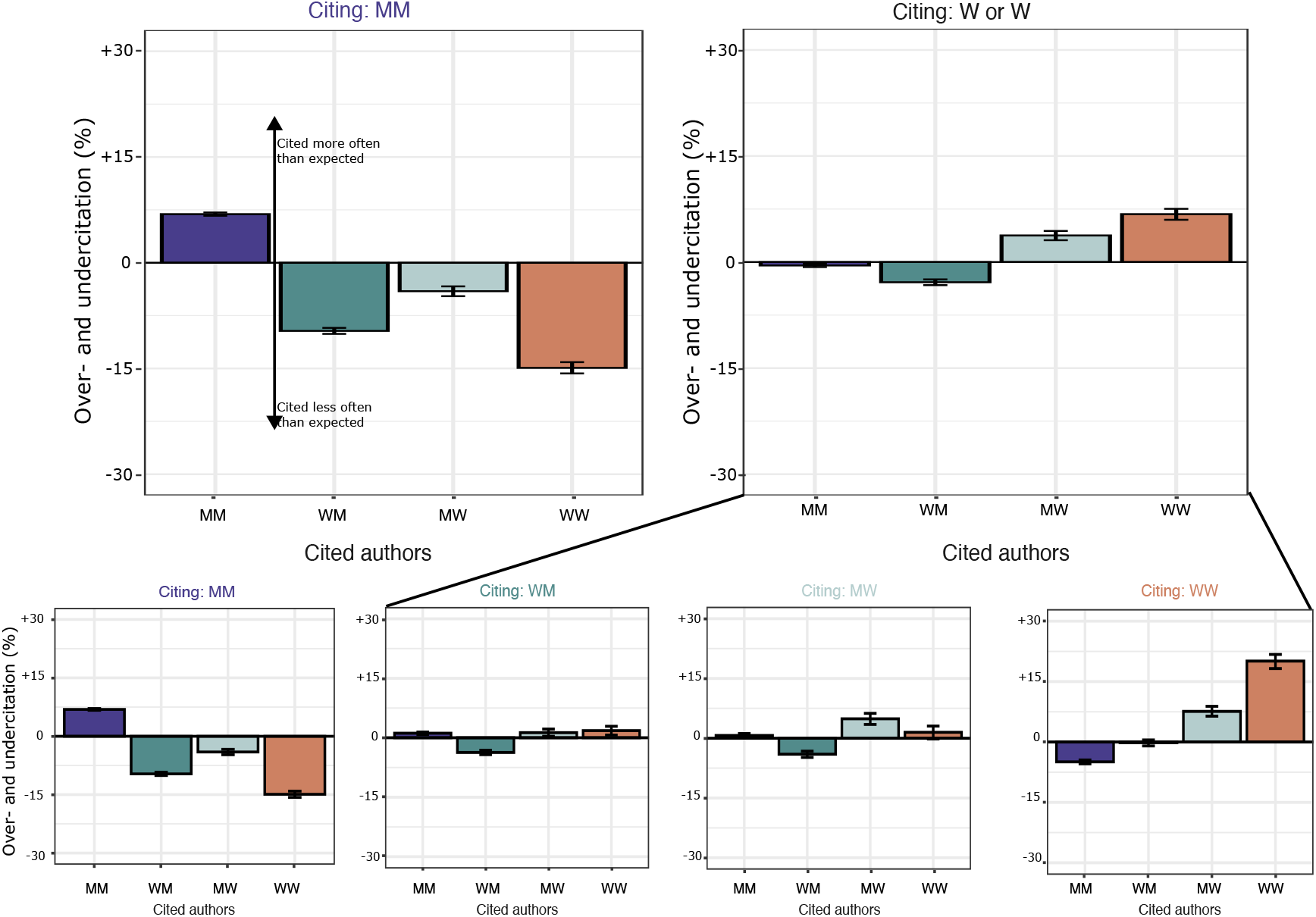
Effect of author gender on citation behavior. Degree of over- and undercitation of different author genders within MM and W∪W reference lists. **Top:** Papers with men as both first and last author overcite men to a greater extent than papers with women as either first or last author. **Bottom:** Full breakdown of gendered citation behavior within MM, WM, MW, and WW reference lists. Bars represent overall over/undercitation. Error bars represent the 95% CI of each over- and undercitation estimate, calculated from 500 bootstrap resampling iterations. Similar breakdowns for author gender within each of the three clusters can be found in the Supplementary Information (Extended Data Figures 7,8,9).

This pattern of results confirmed previous findings that undercitation of women-led papers occurs to a greater extent within men-led reference lists. A key question here is the extent to which this outcome could be driven by other factors. Dworkin and colleagues previously explored the possibility that citation patterns were driven by the overcitation of a few high-impact papers, or by the citation patterns of highly-productive senior authors. However, they found that these factors had little impact on either the extent of citation imbalance or the gender differences in citation behavior (13). Ruling out these alternative explanations suggests that author gender plays a causal role in citation practices.

### Group 1 Papers

Of the 52,112 articles published in journals in this group between 2009 and 2023, 54.6% were written by MM teams and 45.4% were written by W ∪ W teams. After separating citing articles by author gender, we observed that papers led by men first and last authors drove overcitation of men-led papers and undercitation of women-led papers. Specifically, within MM reference lists (*n* = 28, 474 papers), other MM papers were cited 10.6% more than expected (95% CI = (10.3%, 10.9%), one-sample *t* = 62.30, df = 499, *p* < 0.0001), WM papers were cited 14.8% less than expected (95% CI = (−15.5%, -14.2%), one-sample *t* = 44.35, df = 499, *p* < 0.0001), MW papers were cited 10.6% less than expected (95% CI = (−11.8 %, -9.3%), one-sample *t* = 16.72, df = 499, *p* < 0.0001), and WW papers were cited 27.8% less than expected (95% CI = (−29.0%, -26.4%), one-sample *t* = 40.25, df = 499, *p* < 0.0001). Within W ∪ W reference lists (*n* = 23, 638 papers), MM papers were cited 4.1% more than expected (95% CI = (3.7%, 4.5%), one-sample *t* = 20.53, df = 499, *p* < 0.0001), WM papers were cited 7.7% less than expected (95% CI = (−8.4 %, -7.0%), one-sample *t* = 21.71, df = 499, *p* < 0.0001), and WW papers were cited 6.0% less than expected (95% CI = (−7.4%, -4.4%), one-sample *t* = 7.65, df = 499, *p* < 0.0001). We did not find a significant gap for citation of MW papers by W ∪ W teams (one-sample *t* = 1.68, df = 499). Welch two-sample *t*-tests comparing the citation rates between MM- and W ∪ W-led reference lists for each gender category found a significant difference in each case (MM: *t* = 561.59, df = 977, *p* < m.p.; WM: *t* = -324.09, df = 994.11, *p* < m.p.; MW: *t* = -234.35, df = 997.06, *p* < m.p.; WW: *t* = -466.15, df = 983.05, *p* < m.p.), confirming previous findings that undercitation of women-led papers occurs to a greater extent within men-led reference lists within this group.

Within the W ∪ W group, the citation proportions of the WM, MW, and W W subgroups were similar to those found by Ref. (13): WM and MW groups still overcited MM papers and undercited WW papers relative to expectation, but at less than half the rate of MM teams, while WW papers overcited WW teams (at approximately a third of the rate that MM teams overcited MM papers). In addition, the fact that the citation pattern in this group was most similar to the pattern found by Dworkin *et al*. (2020) lends support to the idea that the most prominent journals are best-represented by this pattern (13).

### Group 2 Papers

Of the 80,261 articles published in journals in this group between 2009 and 2023, 38.2% were MM and 61.8% were W ∪ W. After separating citing articles by author gender, we observed that here, the overcitation of women-led papers was driven by women-led teams. Papers from men-led teams (*n* = 30, 660) cited at roughly the expected citation proportions across gender groups. In contrast, within W W reference lists (*n* = 49, 601 papers), MM papers were cited 5.2% less than expected (95% CI = (−5.6 %, -4.9%), one-sample *t* = 28.42, df = 499, *p* < 0.0001), MW papers were cited 6.3% more than expected (95% CI = (5.2 %, 7.4%), one-sample *t* = 11.55, df = 499, *p* < 0.0001), and WW papers were cited 16.2% more than expected (95% CI = (15.0%, 17.4%), one-sample *t* = 26.04, df = 499, *p* < 0.0001). WM papers were cited at roughly the expected proportions. Welch two-sample t-tests comparing the citation rates between MM- and W ∪ W-led reference lists for each gender category found a significant difference in each case (MM: *t* = 448.26, df = 943.41, *p* < m.p.; WM: *t* = -140.58, df = 943.41, *p* < m.p.; MW: *t* = -119.52, df = 947.03, *p* < m.p.; WW: *t* = -404.59, df = 960.08, *p* < m.p.). Within the W ∪ W group, the citation patterns for each subgroup were roughly similar, although WW subgroups tended to overcite WW teams at approximately double the rate of WM and MW teams.

### Group 3 Papers

Of the 51,340 articles published in journals in this group between 2009 and 2023, 44.9% were MM and 55.1% were W ∪ W. After separating citing articles by author gender, we observed that papers led by men first and last authors once again largely accounted for the overcitation of men-led papers and undercitation of women-led papers. Specifically, within MM reference lists (*n* = 23, 035 papers), other MM papers were cited 6.6% more than expected (95% CI = (6.1%, 7.1%), one-sample *t* = 27.98, df = 499, *p* < 0.0001), WM papers were cited 10.3% less than expected (95% CI = (−11.1%, -9.6%), one-sample *t* = 25.57, df = 499, *p* < 0.0001), and WW papers were cited 13.6% less than expected (95% CI = (−15.2%, -12.3%), one-sample *t* = 18.50, df = 499, *p* < 0.0001). MW papers were cited roughly as often as expected. Within W ∪ W reference lists (*n* = 28, 305 papers), WM papers were cited 4.1 % less than expected (95% CI = (−4.8 %, -3.4%), one-sample *t* = 11.30, df = 499, *p* < 0.0001), MW papers were cited 5.1% more than expected (95% CI = (3.6 %, 6.4%), one-sample *t* = 7.63, df = 499, *p* < 0.0001), and WW papers were cited 3.3% more than expected (95% CI = (1.6%, 4.9%), one-sample *t* = 4.19, df = 499, *p* < 0.0001). MM papers were cited roughly as often as expected. Welch two-sample *t*-tests comparing the citation rates between MM- and W ∪ W-led reference lists for each gender category found a significant difference in each case (MM: *t* = 430.29, df = 991.89, *p* < m.p.; WM: *t* = -255.22, df = 988, *p* < m.p.; MW: *t* = 116.87, df = 979.89, *p* < m.p.; WW: *t* =351.73, df = 994.93, *p* < m.p.), confirming previous findings that undercitation of women-led papers occurs to a greater extent within men-led reference lists.

However, when we analyzed each gender category separately, we once again found a link between the increased citation of women-led work and the increased leadership role of women on the citing team. In this group, WM and MW teams both cited at roughly the expected rates (slightly underciting WM papers), but WW teams overcited WW papers at roughly the same rate that they were undercited by MM groups.

Taken together, these findings, summarized in Fig. 5, show that more gender-diverse author teams seem to engage in less biased citing practices across all groups. Teams composed of authors of the same gender were more likely to exhibit a gender bias (homophily) in their citation practices. However, homophily that favored men was more present in the top journals, while homophily driven by women-led teams was more present in lower-tier journals.

**Figure 5.**
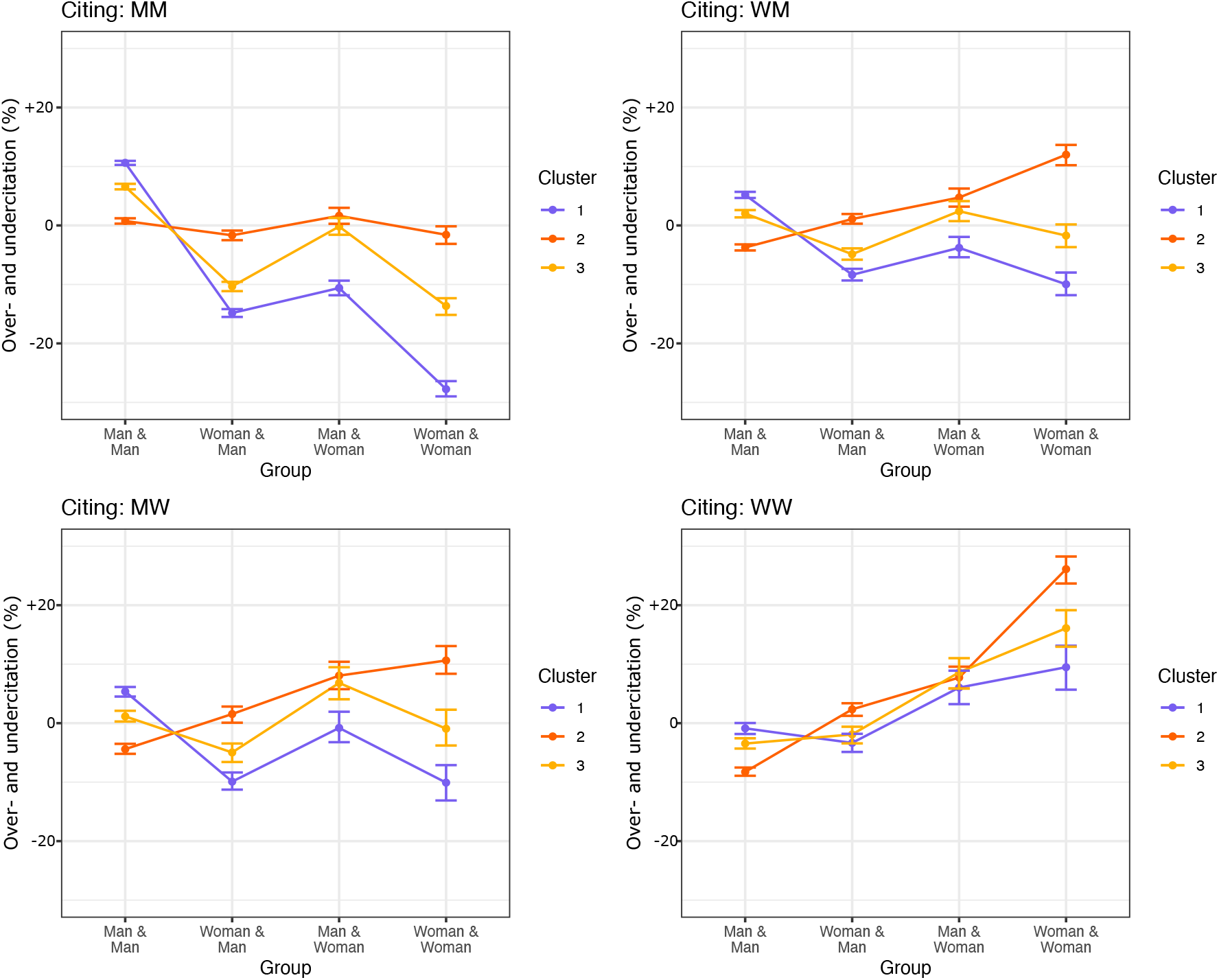
Effect of author gender on citation behavior. Degree of over- and undercitation of different author genders within MM and W∪W reference lists, for each group. Error bars represent the 95% CI of each over- and undercitation estimate, calculated from 500 bootstrap resampling iterations.

## Discussion

In this study, we sought to determine whether gender imbalance in neuroscience citations is unique to the most prestigious journals or extends to the field as a whole. In a significantly larger database than utilized in previous studies, we confirmed the presence of a gender gap in citations. Notably, the magnitude of the gap was somewhat smaller in this wider dataset, and its partial diminution was due to the interaction of three separate subgroups: one group that, as reported earlier, overcites MM papers and undercites all other categories; one group that overcites papers with a woman as a last author; and one group that undercites papers with a woman as a first author. These groups are not distributed evenly across the field; more prestigious journals tend to have articles with citation gaps favoring men, while less prestigious journals tend to have articles with citation gaps favoring women.

### Potential Drivers of Homophilic Behaviors

As demonstrated in this work, papers with single-gender author teams were more likely to have gender gaps than papers with a man and a woman as first or last author. These gaps generally favored teams with the same gender makeup as the citing authors, and undercited teams with the opposite gender makeup. This behavior is known as homophily, and is consistent with other studies of gender citation gaps reporting similar homophily (31–33).

Previous studies have suggested that gender homophily in citation patterns can arise due to gender homophily in professional pathways (33) and associated research topics (31, 33). This suggestion has been motivated by prior work demonstrating that some measures of identity correlate with research topic (33, 34). In our study, an inspection of paper titles provides some qualitative (but not quantitative) evidence for topic variation across groups (Extended Data Figures 10,11,12): for example, Group 2 seems to emphasize wet-lab and bench science, while Group 3 seems to have more of a focus on translational and human-centric studies. However, our statistical model accounts for the gender proportions in the field as a whole, and prior work has demonstrated that the gender citation gap persists even when accounting for sub-field variations in gender proportions (13), suggesting that other explanations are needed.

Gender homophily in citation patterns can arise from a range of psychological and social factors. Some are more proximal, impacting the moment of citation directly, while others are more distal, impinging on citations indirectly. An example of a proximal factor is cognitive bias: explicit or implicit bias can lead a person to favor colleagues of the same gender in different areas of scientific practice. For citation practices in particular, implicit bias against marginalized groups (e.g., women and people of color) and in favor of majority groups (e.g., men and white people) has been suggested as a primary driver (13, 20, 28, 35). An example of a distal factor is a scholar’s awareness of the work of others from the same or different identity groups. Awareness can be influenced by visibility: men’s greater visibility over women’s in science, due to structural inequality, may lead the average scholar to be less familiar with the work of women and more familiar with the work of men. This difference in familiarity could explain why the average scholar might think to cite the (known) work of men (13, 20, 35).

At an intermediate scale between proximal bias and distal awareness, gendered homophily in citation patterns could arise from the gender proportions of collaborative relationships (36–38). Prior work has shown that the overrepresentation of men in the co-authorship networks of men-led teams explains significant variance in the overcitation of MM papers (13). Some women may also have overrepresentation of other women in their co-authorship networks, either because they (i) choose to collaborate with other women to boost confidence and camaraderie in a field dominated by men, or (ii) the exclusion (or expectation of exclusion) of women from men-led teams leads them to preferentially work with other women (37, 38). When an author from a given gender group has an overrepresentation of their own gender in coauthorship or collaboration networks, that imbalance can enhance the author’s awareness of work from the same gender. Because awareness of a scholar’s work can influence the probability of citing them, gender representation in collaboration networks is a potential mechanism for gender homophily in citation patterns.

Factors such as awareness, collaboration, and implicit bias are candidates for passive drivers of citation imbalances. Could active drivers be at play as well? Prior work has described the phenomenon of ‘activist choice homophily’, where members of a minority population band together against shared structural barriers to help each other overcome these barriers (20, 39), or even more simply a conscious attempt to close the gap of overcitation of men by collaboration. This may be a partial explanation for the behaviors we observe in Group 2, where overcitation of groups with a woman as the last author are driven primarily by women-led groups. The last author is generally the more senior author, and would likely be more connected to the network of other women in neuroscience. Since teams with a woman as the last author are overcited, rather than just WW groups or those with any woman as an author, this suggests a specific effect associated with the last author. This pattern may indicate an intentional practice among networks of women PIs to highlight the contributions of other women, consistent with a distributive justice model of ethics (40).

### Differential Impact of Gender Imbalances

Although Group 1 manifested a gender citation balance in favor of men and Group 2 manifested a gender citation balance in favor of women, the former behavior has a markedly more negative impact on the field than the latter. First, a citation imbalance in favor of men hurts the minority group—women—who are already disadvantaged in terms of compensation (1), hiring and promotions (2–4), authorship (5), teaching evaluations (6, 7), grant funding (8, 9), collaboration credit (10), publication quantity (11), and tier of journal publication (12). Second, a citation imbalance in the high impact journals means a markedly larger citation decrement for the undercited group than a citation balance in the low impact journals. Because the high-impact journal citation imbalance favors men, this behavior produces a significant citation decrement to women, directly harming career advancement. So long as these journals have an outsize effect on the direction of the field, any inequities in citation practices of authors publishing in those journals will have outsize effects on authors’ careers as well.

### Considerations for Mentors

The data reported here can inform individual choices regarding citation practice, as well as factors to consider in building teams and mentoring trainees. In building teams, mixed author groups may be able to combine enhanced awareness of both man-authors papers and woman-authored papers, given homophilic tendencies in the field. We see some evidence for this possibility in our data. Man-led homophily was highest in high-status Group 1, where MM papers made up more than half of the pool, and was lowest in low-status Group 2, where MM papers accounted for only 40 percent of the total. Woman-led homophily was highest in low-status Group 2, where papers with at least one woman accounted for approximately 60 percent of the papers, and lowest in Group 1, where there were far fewer women. Papers with both a man and a woman as an author (in either order) tended to overcite MM papers more in Group 1, and overcite papers with women as last author more in Group 2, showing a tendency for non-homogeneous author teams to default towards the most prominent pattern in a group. However, in both cases, this behavior was not as pronounced as it was by the most prominent gender group (i.e., in Group 1, MW and WM groups overcited MM papers to a lesser extent than MM authors did). In Group 3, where both homophilous patterns (MM overcitation by men, and _W overcitation by women) were present, non-homogeneous author teams cited closer to the MM pattern, but to a far lesser degree. This pattern suggests that the collaborations of researchers of multiple genders could potentially help to close the gender gaps in STEM—a point supported by prior work (38). This finding adds to the body of existing evidence showing the utility of diverse teams in science (31, 34, 41).

Beyond building teams, the data reported here can also inform how more senior scholars mentor trainees, and particularly women trainees. We observe that despite the counteracting homophilic citation preferences of each group, the undercitation of WM papers persists across the field at large (5.8%). This gap appears to be driven by articles in Group 1, and to a lesser extent, Group 3. In these groups, man-led homophily leads to the undercitation of all articles with a woman as an author (WM, MW, and WW). While the overcitation of women as a last author in Group 2 counters this gap for WW and MW articles, there is no such counterbalance for WM articles and therefore the gap persists. This pattern of findings suggests that women first authors (and men, though there are other forces assisting them) may lack access to the same networks of collaboration as students working for a woman as a PI. All trainees should have the benefit of PIs who devote time and energy to building community, resources, and networks for and with women scholars.

## Methodological Considerations and Limitations

As noted earlier, this work is subject to several limitations. First, the methods used for gender determination are limited to binary man and woman gender assignments. This study design, therefore, cannot properly encapsulate intersex, transgender, and/or nonbinary identities. Second, this study investigates inequalities solely along gender lines. Future work could involve a more intersectional examination, including factors such as race, class, sexuality, disability and citizenship.

## Acknowledgments

The content is solely the responsibility of the authors and does not necessarily represent the official views of any of the funding agencies.

## Contributions

Conceptualization: K.H. & D.S.B.; methodology: K.H. & E.G.T.; data curation: K.H.; formal analysis: K.H.; writing: K.H. (original draft) and K.H. & D.S.B. (review and editing); funding acquisition: K.H. & D.S.B.; supervision: D.S.B.

## Data availability

Data and materials for this study were downloaded from Web of Science.

## Code availability

The code used for sampling, processing and analysis can be accessed at https://github.com/jdwor/gendercitation.

## Methods

### Data collection

Data for this study were drawn from the Web of Science (WoS) database, which indexes journals according to the Science Citation Index Expanded. We selected the neuroscience journals with the fifty highest Eigenfactor scores. Eigenfactor scores provide metrics of journal prominence and influence within a given field by giving a count of incoming citations, weighted by the impact of the citing journal (27). A full list of the journals cited can be found in the Supplementary Information (Table 1).

From these 50 journals, we downloaded data from all articles published between 1995 and 2023. We included all articles classified as articles, review articles, or proceedings papers labelled with a digital object identifier (DOI) in our analyses. For each article, we downloaded the author names, reference lists, publication date, and DOI, and we matched each paper’s cited references to the other papers included in the dataset through their DOIs.

Because papers before 2006 were less likely to include authors’ first names, we used CrossRef API (www.crossref.org) to try to identify author first names on papers published before that time. We then implemented a name disambiguation algorithm (see Dworkin *et al*., 2020 (13) for details) to minimize the number of papers for which we only had access to authors’ initials. After we disambiguated authors, we used the methods described in Ref. (13) to predict author gender, remove self-citations, and develop a citation network. Although more details can be found in our past work, we have provided a brief summary below.

### Author gender determination

For authors with available first names, gender was assigned to first names using a database created from the ‘gender’ package in R with the Social Security Administration (SSA) baby name dataset. For names that were not included in the SSA dataset, gender was assigned using Gender API (http://gender-api.com/), a paid service that supports roughly 6,000,000 unique first names across 191 countries. We assigned ‘man’ (‘woman’) to each author if their name had a probability ≥0.70 of belonging to someone labeled as ‘man’ (‘woman’) according to a given source (42). In the SSA dataset, man and woman labels correspond to the sex assigned to children at birth; in the Gender API dataset, man and woman labels correspond to a combination of sex assigned to children at birth and genders detected in social media profiles.

Gender could be assigned to both the first and last author of 77.7% of the papers in the dataset. For the remaining 22.3% gender data was either unavailable, or the gender was not certain. Approximately 14.6% of unique first names had an uncertain gender, and an additional 13.2% were unavailable. We performed the following analyses using the articles for which gender could be assigned with high probability to both authors (*n* = 281, 776). ^3^

### Interpretation of author gender assignments

As in previous papers (13, 20, 28), we use the term ‘gender’ in our analysis not to directly refer to the sex of the author, as assigned at birth or chosen later, nor to directly refer to the gender of the author, as socially-assigned or self-chosen, but instead to model the probability that an author has a certain gender based on their name. Specifically, ‘woman’ refers to an author with a name with a probability ≥0.70 of being given to a child assigned female at birth or someone identifying as a woman on social media; ‘man’ likewise refers to an author with a name with a probability ≥0.70 of being given to a child assigned male at birth or someone identifying as a man on social media. The author’s actual sex or gender is not identified.

Given the limitations of this probabilistic method, the predicted gender for any author may in fact be quite different from their true sex or gender. Authors may also be intersex, transgender, or nonbinary, none of which is accounted for in this study. However, in many cases, citers will also not know the true sex and/or gender of the authors they cite, and may infer either based on the author’s name. In this way, our probabilistic analysis nontrivially captures citation biases arising from both known and inferred gender.

### Removal of self-citations

For this study, self-citations were removed from all analyses of gendered citation behavior. Although self-citations themselves have been found to have relevant gendered properties (13, 43), their removal in this study allowed us to isolate more comparable external citation behaviors of men and women in the field. We defined self-citations as papers for which either the cited first or last author was the first or last author on the citing paper. While this definition is somewhat restrictive, it is the only type of self-citation for which the author gender of the cited paper is necessarily determined by the author gender of the citing paper. In addition, in our previous work (13), we demonstrated that using broader definitions of self-citation that included the entire author list of the cited or citing paper had little to no impact on the results.

We further explored the relative presence of self-citations in this data, across author genders and across clusters, as seen in the Supplementary Information (Table 2 and Extended Data Figures 3,4,5,6).

### Statistical Analyses

Many of the analyses in this study compared the observed citation behavior with the rates at which MM, WM, MW, and WW papers would be expected to appear in reference lists if gender were irrelevant. To obtain expected rates after controlling for various characteristics that might be correlated with gender, we fit a Generalized Additive Model (GAM) on the category label MM, WM, MW, and WW, considering the following features: (1) month and year of publication; (2) combined number of publications by first and last author; (3) number of total authors on the paper; (4) journal of publication; and (5) whether the paper was a review. We applied this model to each paper to obtain the probability that the paper belonged in each of the four categories. By summing the probabilities for each of the papers cited in an article, we were able to calculate the rate at which the different gender categories might be expected to appear in that article’s reference list if author gender were independent of citation rates and dependent on the other characteristics in the model. The GAM was fit using the mgcv package in R (44), using penalized thin-plate regression splines for estimating smooth terms of publication date, author experience and team size. For the primary analyses, univariate thin-plate splines were used for the smooth terms, and no interactions between variables were included in the model.

We present all estimates in this study with a confidence interval (CI), a *p*-value, or both. We calculated the CIs by bootstrapping citing papers, or randomly sampling citing papers with replacement. This procedure maintained the dependence structures of the clusters of cited articles within citing articles. The null model used to obtain *p*-values was that there was no over- or undercitation:

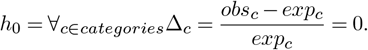

Because statistical significance is assessed for multiple primary comparisons, all presented *p*-values were corrected according to the Tukey method.

In the following sections, we describe the formal statistical analyses that we used to address our distinct hypotheses. In each subsection, we state the hypothesis first, followed by the analysis used to test it. All hypotheses were tested for the set of articles published between 2009 and 2023 for which both first and last author gender could be predicted.

### Hypothesis 1: The overall citation rate of women-led papers will be lower than expected given relevant characteristics of papers

To test this hypothesis, we first estimated the expected number of citations given to each author gender category. We calculated this expectation by summing over the GAM-estimated probabilities for all papers contained within the reference lists of citing papers. These totals, therefore, reflect the expected number of citations given to MM, WM, MW, and WW papers if author gender was conditionally independent of citation behavior, given the paper characteristics included in the model described above.

To calculate the observed number of citations given to each group, we simply summed over the {MM, WM, MW, WW} variable for all of the papers contained within the reference lists of papers published between 2009 and 2023. These values were compared by calculating the percent difference from expectation for each author gender group. For example, for WW papers, this percent change in citation would be defined as,

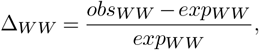

where *obs*_*W W*_ is the number of citations given to WW papers between 2009 and 2023, and *exp*_*W W*_ is the expected number of citations given to WW papers between 2009 and 2023.

Notably, performing the summation over all citations results in the upweighting of articles with many citations and the downweighting of articles with few citations. This approach helps to improve the stability of the estimates. In addition, this approach does not appear to be sensitive to high-influence observations: our previous work (13) found little difference between this method versus using the mean of article-level over/undercitation.

### Hypothesis 2: Undercitation of women-led papers will occur to a greater extent within the most influential journals

To test this hypothesis, we used very similar metrics to those described in the previous section. The primary difference is that instead of calculating the observed and expected citations by summing over the citations within all reference lists between 2009 and 2023, in this section we performed those summations separately for reference lists in papers published in the five journals with the highest Eigenfactor, and papers published in the remaining 45 journals. For example, to estimate the over/undercitation of WW papers within the reference lists of papers in the top five journals, we define,

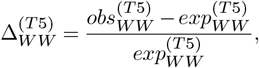

where 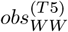 is the total number of citations given to WW papers within the reference lists of the top five journals, and *WW* 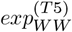 is the expected number of citations given to WW papers within these reference lists.

### Hypothesis 3: Undercitation of women-led papers will occur to a greater extent within men-led reference lists

To test this hypothesis, we used very similar metrics to those described in the previous section. The primary difference is that instead of calculating the observed and expected citations by summing over the citations within all reference lists between 2009 and 2023, in this section we performed those summations separately for reference lists in papers with men as first and last author (MM papers) and papers with women as first or last author (W ∪ W papers). For example, to estimate the over/undercitation of WW papers within the reference lists of MM papers, we define,

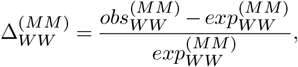

where 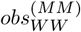 is the total number of citations given to WW papers within MM reference lists, and 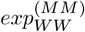 is the expected number of citations given to WW papers within MM reference lists.

We performed this test not only within the entire corpus, but also within each of our three groups.

## Supplementary Information

### Journals, Eigenfactors, and Group Membership

This table includes journal name and Eigenfactor as of September 2023, for the fifty journals included in the analysis. It also includes group membership, determined as described elsewhere.

**Table 1.** Journal names, eigenfactor, and group membership, sorted by Eigenfactor as of September 2023.

| Journal Name | Eigenfactor | Group |  |  |
| --- | --- | --- | --- | --- |
|  |  | 1 | 2 | 3 |
| *Neuron* | 0.1293 | X |  |  |
| *Nature Neuroscience* | 0.1044 | X |  |  |
| *Neuroimage* | 0.0988 | X |  |  |
| *Journal of Neuroscience* | 0.0793 | X |  |  |
| *Frontiers in Neuroscience* | 0.0668 |  |  | X |
| *Frontiers in Neurology* | 0.0642 |  |  | X |
| *Brain* | 0.0556 |  |  | X |
| *Nature Human Behavior* | 0.0528 | X |  |  |
| *Molecular Psychiatry* | 0.0511 |  |  | X |
| *Neuroscience and Biobehavioral Reviews* | 0.0449 |  | X |  |
| *Journal of Alzheimers Disease* | 0.0400 |  | X |  |
| *Nature Reviews Neuroscience* | 0.0393 |  |  | X |
| *Brain Behavior and Immunity* | 0.0358 |  | X |  |
| *Cerebral Cortex* | 0.0358 |  |  | X |
| *Biological Psychiatry* | 0.0342 |  | X |  |
| *Annals of Neurology* | 0.0335 |  |  | X |
| *Human Brain Mapping* | 0.0318 |  |  | X |
| *Trends in Cognitive Sciences* | 0.0312 |  |  | X |
| *Frontiers in Cellular Neuroscience* | 0.0310 |  | X |  |
| *Neuropsychopharmacology* | 0.0307 |  | X |  |
| *Frontiers in Aging Neuroscience* | 0.0285 |  | X |  |
| *Acta Neuropathologica* | 0.0282 |  | X |  |
| *Journal of Neuroinflammation* | 0.0276 |  | X |  |
| *Pain* | 0.0260 |  | X |  |
| *Journal of Physiology-London* | 0.0251 | X |  |  |
| *Molecular Neurobiology* | 0.0248 |  | X |  |
| *Frontiers in Human Neuroscience* | 0.0244 |  |  | X |
| *Neural Networks* | 0.0239 | X |  |  |
| *Neuroscience* | 0.0233 |  | X |  |
| *Brain Sciences* | 0.0226 |  | X |  |
| *Frontiers in Molecular Neuroscience* | 0.0211 |  | X |  |
| *Neuropharmacology* | 0.0210 |  | X |  |
| *European Journal of Neurology* | 0.0203 |  |  | X |
| *Sleep* | 0.0203 |  |  | X |
| *Current Opinion in Neurobiology* | 0.0201 | X |  |  |
| *Neuroscience Letters* | 0.0195 |  | X |  |
| *Neurobiology of Aging* | 0.0182 |  | X |  |
| *Neurobiology of Disease* | 0.0180 |  | X |  |
| *Behavioural Brain Research* | 0.0179 |  | X |  |
| *eNeuro* | 0.0179 | X |  |  |
| *Journal of Cerebral Blood Flow and Metabolism* | 0.0177 | X |  |  |
| *Trends in Neurosciences* | 0.0175 |  |  | X |
| *Psychoneuroendocrinology* | 0.0174 |  | X |  |
| *Cortex* | 0.0173 |  | X |  |
| *Journal of Neurotrauma* | 0.0172 |  |  | X |
| *Acta Neuropathologica Communications* | 0.0170 |  | X |  |
| *Journal of Neurophysiology* | 0.0169 | X |  |  |
| *Journal of Stroke & Cerebrovascular Diseases* | 0.0166 | X |  |  |
| *ACS Chemical Neuroscience* | 0.0162 |  |  | X |
| *Multiple Sclerosis Journal* | 0.0160 |  |  | X |

### Sensitivity to start year of citing paper pool

For all analyses reported in the primary manuscript, the pool of citing papers included papers published between 2009 and 2023. The 2009 start date was used to ensure consistency with past work. However, to ensure that there was nothing unique about 2009 as a starting year, we calculated the citation gaps for pools that began in neighboring years, i.e. in 2008 and 2010, and observed that there was no significant difference.

**Extended Data Figure 1.**
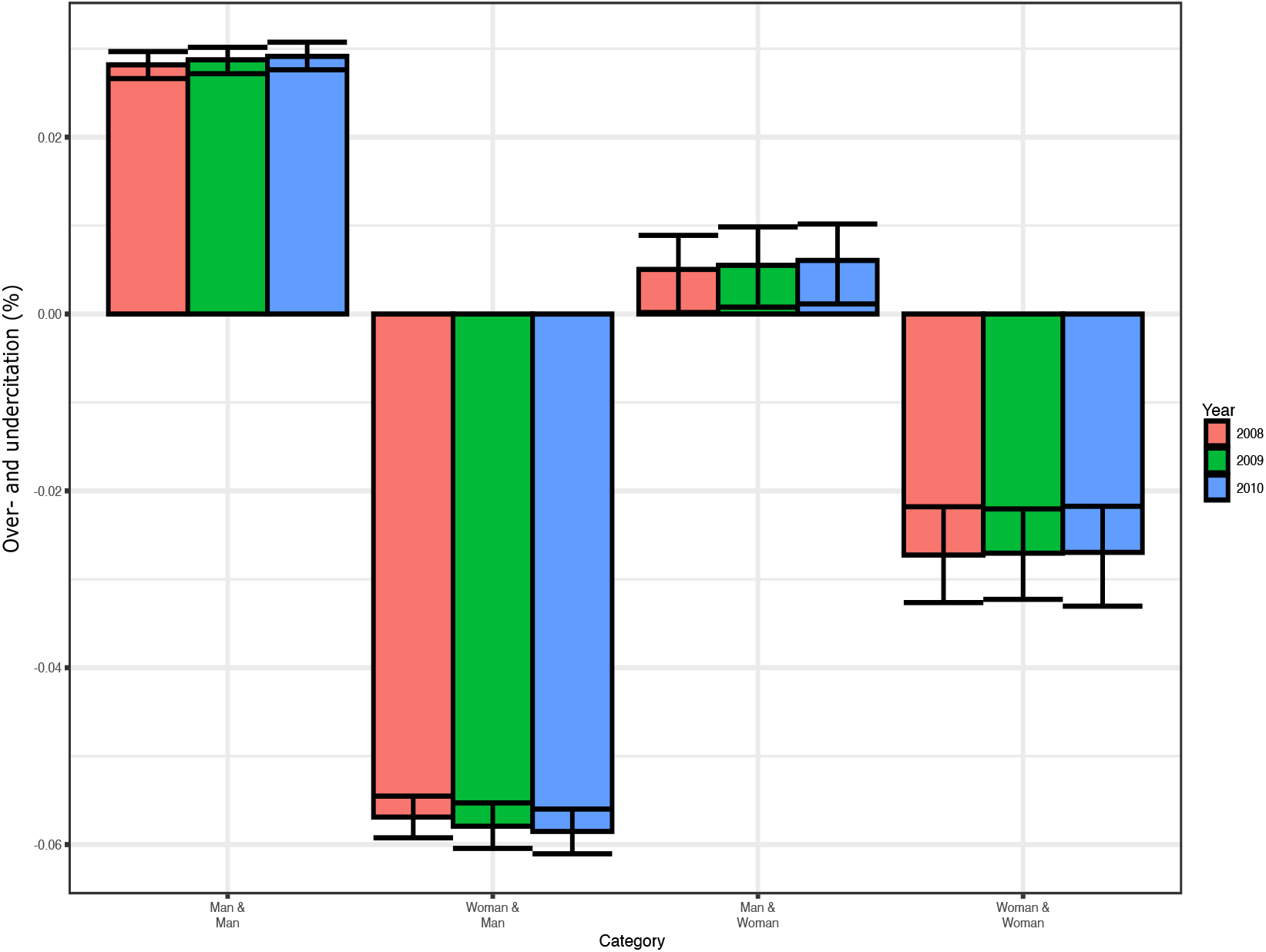
Citation gap across gender categories with different cutoffs for citing pool.

### Rate of change of publishing composition

In our paper, we note the average rate of increase in publication of papers with at least one woman author. We also calculated this rate for each individual journal, separately. We calculated the percent of articles with at least one woman author each year, then regressed this value as a function of the year. 95% confidence intervals are shown. As can be seen, there is marked variability by journal, and at least six of the fifty journals do not display a clear increase in articles with a woman author over time. In addition, three of the top five journals (*Journal of Neuroscience, Nature Neuroscience*, and *Neuron*) have rates of change that are noticeably lower than much of the rest of the field. (There is not a significant relationship between Eigenfactor and rate of increase, however.) The journals with the highest Eigenfactor as of 2023 are highlighted in red.

**Extended Data Figure 2.**
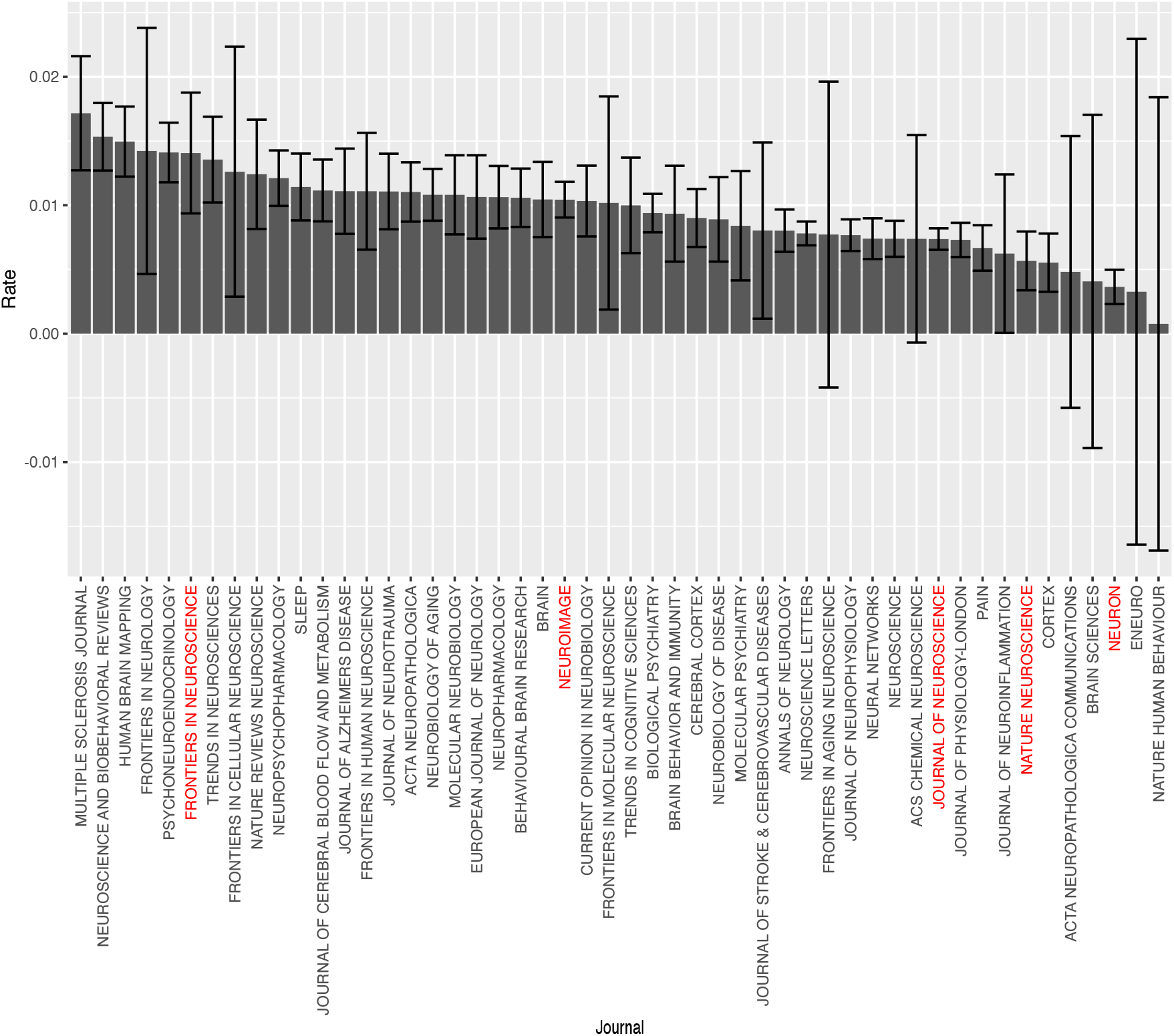
Journal-specific yearly increase in articles with at least one woman as an author.

### Selection of group size and members

The groups described in the primary manuscript were generated by the application of a *K*-means clustering approach with *K* = 3. We chose this number heuristically based not only on elbow plots but because increasing the number of clusters tended to increase the likelihood of singleton clusters. In addition, clustering with three clusters was remarkably stable. After removal of one outlying journal (*Journal of Alzheimer’s Disease*, which had a particularly high rate of MW overcitation), the clustering came out identically across 1000 trials, each with random seeds. (We fit the model on the 49 non-outlying journals, then assigned the *Journal of Alzheimer’s Disease* to the group with the nearest centroid.) By comparison, clustering with additional groups yielded far more variation in cluster sizes. We also verified these clusters using R’s *fanny* method, which returned the same results. We then assigned the outlying journal to the cluster it was closest to.

### Self-citation rates

For all analyses reported in the primary manuscript, self-citations were removed from consideration. While our previous work found that the inclusion of self-citations did not significantly impact patterns of over- and undercitation (13), we did report that papers with a man as last author tended to self-cite at higher rates than teams with a woman as last author.

Given that the journals used in our previous study were not entirely representative, we once again examined potential gendered differences in the rate of self-citations across author genders. We found that, as a proportion of reference list length, MM and WM teams tended to self-cite at higher rates than MW and WW teams. However, as a proportion of self-citations (i.e., as compared to the count of authors’ previous citable papers), rates of self-citation were relatively similar, though slightly higher in WW and MW teams. This is explained in part by the fact that on average, WW and WM teams had less previous citeable papers (23.6 for MM (95% CI 23.5, 23.8), 20.9 for WM (95% CI 20.7, 21.1), 18.4 for MW (95% CI 18.2, 18.6), and 14.2 for WW (95% CI 14.0, 14.4).

We then explored whether self-citation rates varied as a function of group and gender. We found that the pattern of more self-citation in MM and WM papers, and less in MW and WW papers, held true across each group. However, in general, Group 1 had the most self-citation as a function of reference list length, followed by Group 2, and then Group 3.

Finally, we calculated self-citation rates for each individual journal.

**Table 2.** Comparison of self-citation rates across author gender categories, relative to both the length of reference lists and the number of potential self-citations. Confidence intervals are calculated from 1,000 bootstrap resampling iterations.

| Effect | MM teams |  | WM teams |  | MW teams |  | WW teams |  |
| --- | --- | --- | --- | --- | --- | --- | --- | --- |
|  | Est. | 95% CI | Est. | 95% CI | Est. | 95% CI | Est. | 95% CI |
| Mean self-citations by reference list length | 9.3% | [9.2, 9.4] | 8.9% | [8.8, 9.0] | 8.3% | [8.1, 8.5] | 7.2% | [7.0, 7.3] |
| Median self-citations by reference list length | 4.5% | [0, 45.5] | 4.3% | [0, 42.9] | 3.3% | [0, 42.1] | 0.0% | [0, 38.5] |
| Mean self-citations by num. previous papers | 8.1% | [8.0, 8.1] | 8.2% | [8.1, 8.3] | 8.6% | [8.4, 8.7] | 8.5% | [8.4, 8.7] |
| Median self-citations by num. previous papers | 3.6% | [0, 39.1] | 3.6% | [0.0, 40.0] | 3.2% | [0.0, 40.0] | 0.0% | [0.0, 42.9] |

**Extended Data Figure 3.**
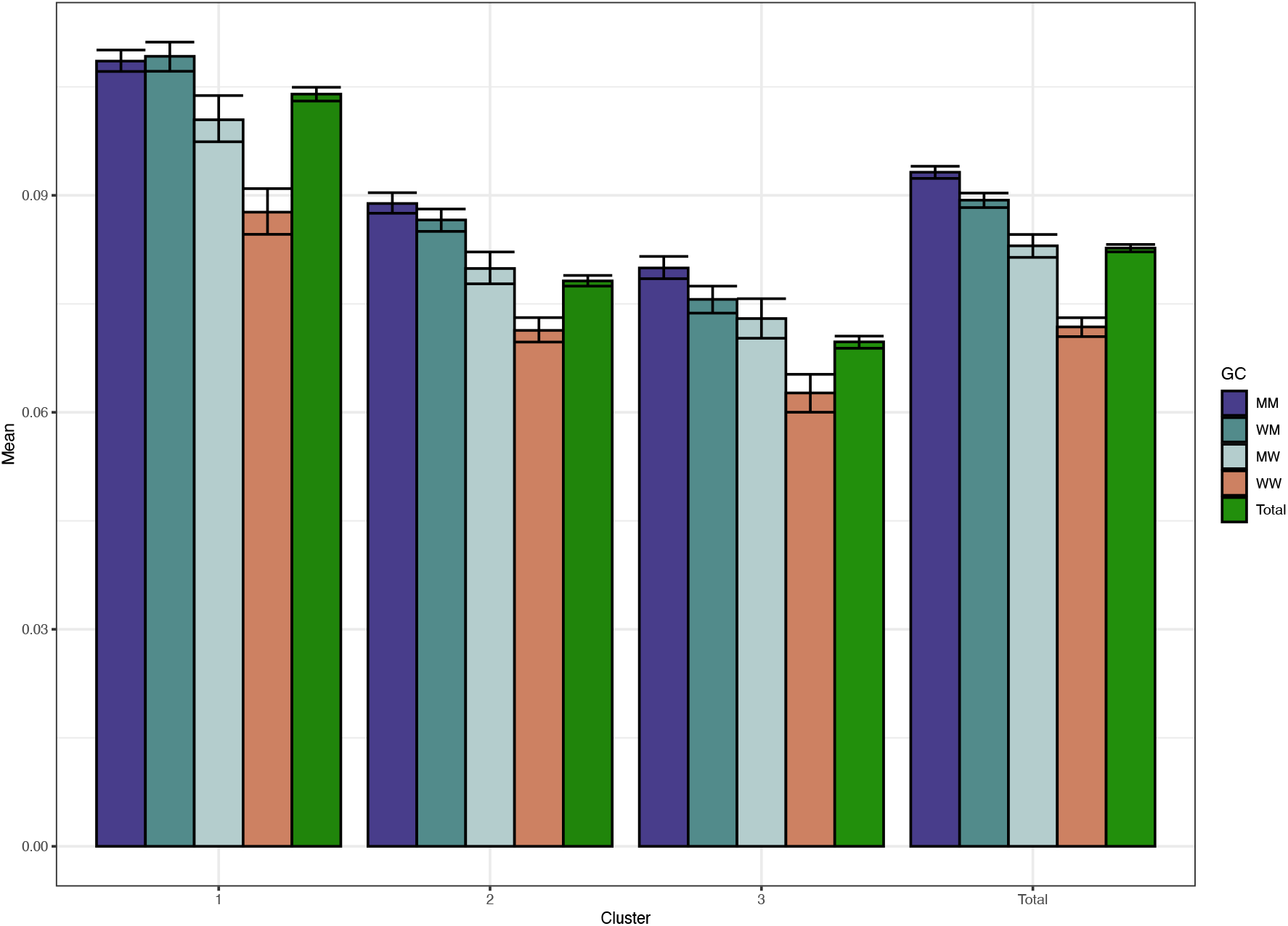
Self-citation rate by gender category, within cluster. Confidence intervals are calculated from 1,000 bootstrap resampling iterations.

**Extended Data Figure 4.**
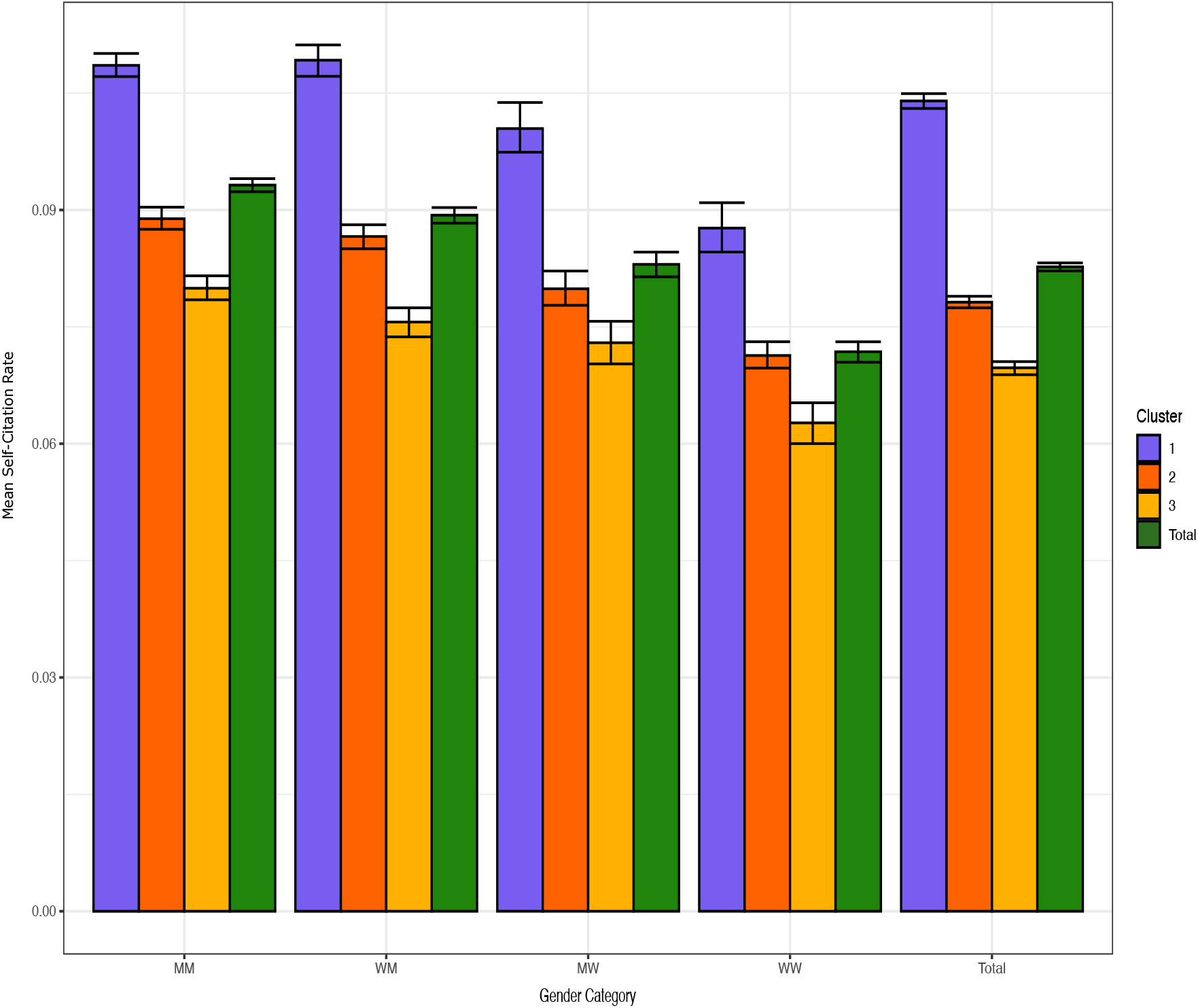
Self-citation rate within group, by gender category. Confidence intervals are calculated from 1,000 bootstrap resampling iterations.

**Extended Data Figure 5.**
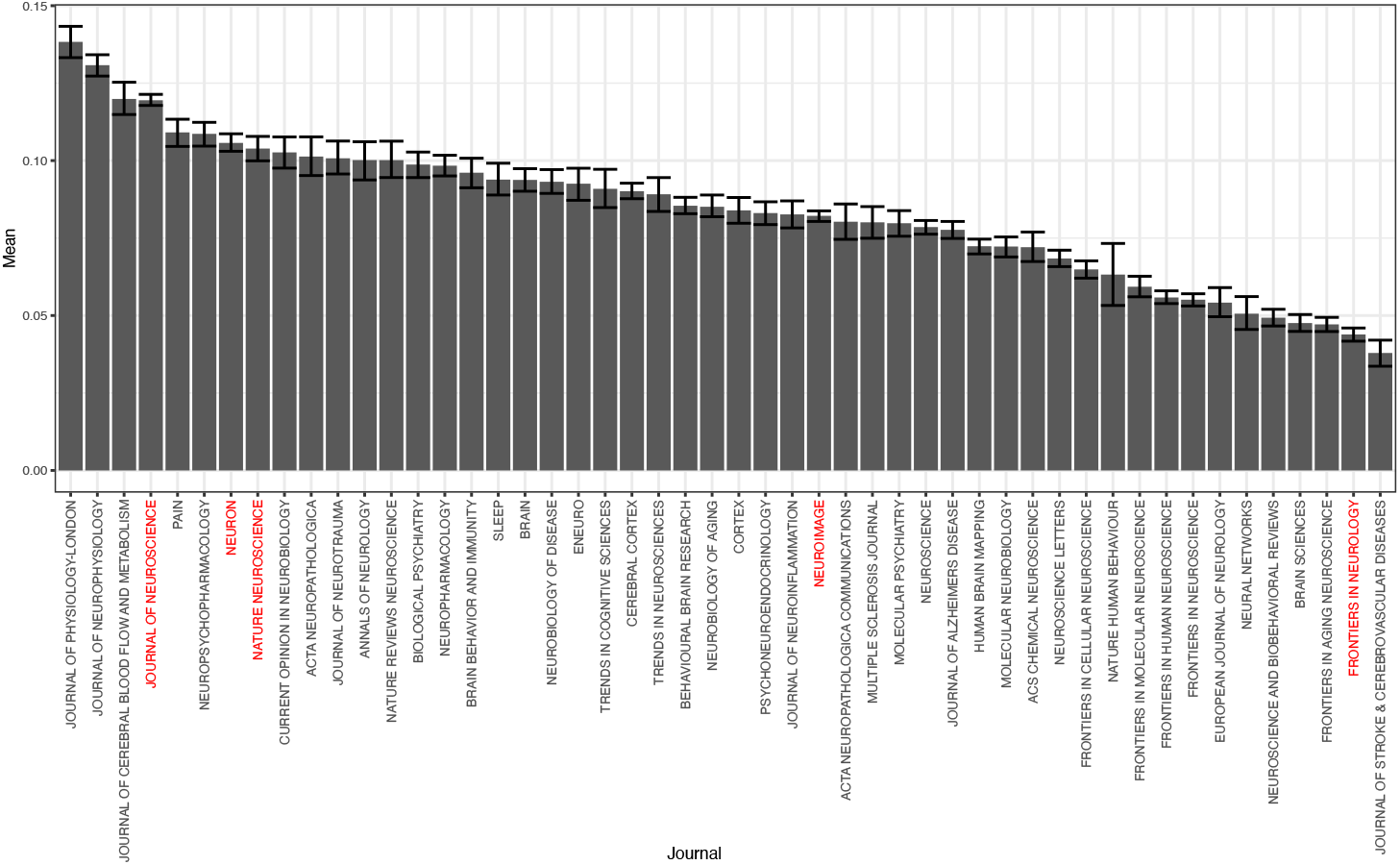
Self-citation rate by journal. Confidence intervals are calculated from 1,000 bootstrap resampling iterations.

### Journal Insularity

We also explored self-citation at the journal level by measuring the proportion of citations made by papers published in a journal to other papers published in the same journal. As shown in the figure below, insularity varied a great deal by journal. *Neural Networks* and *Sleep* seemed to be the most insular and specialized, while journals that published reviews were far more likely to cite mainly other journals.

**Extended Data Figure 6.**
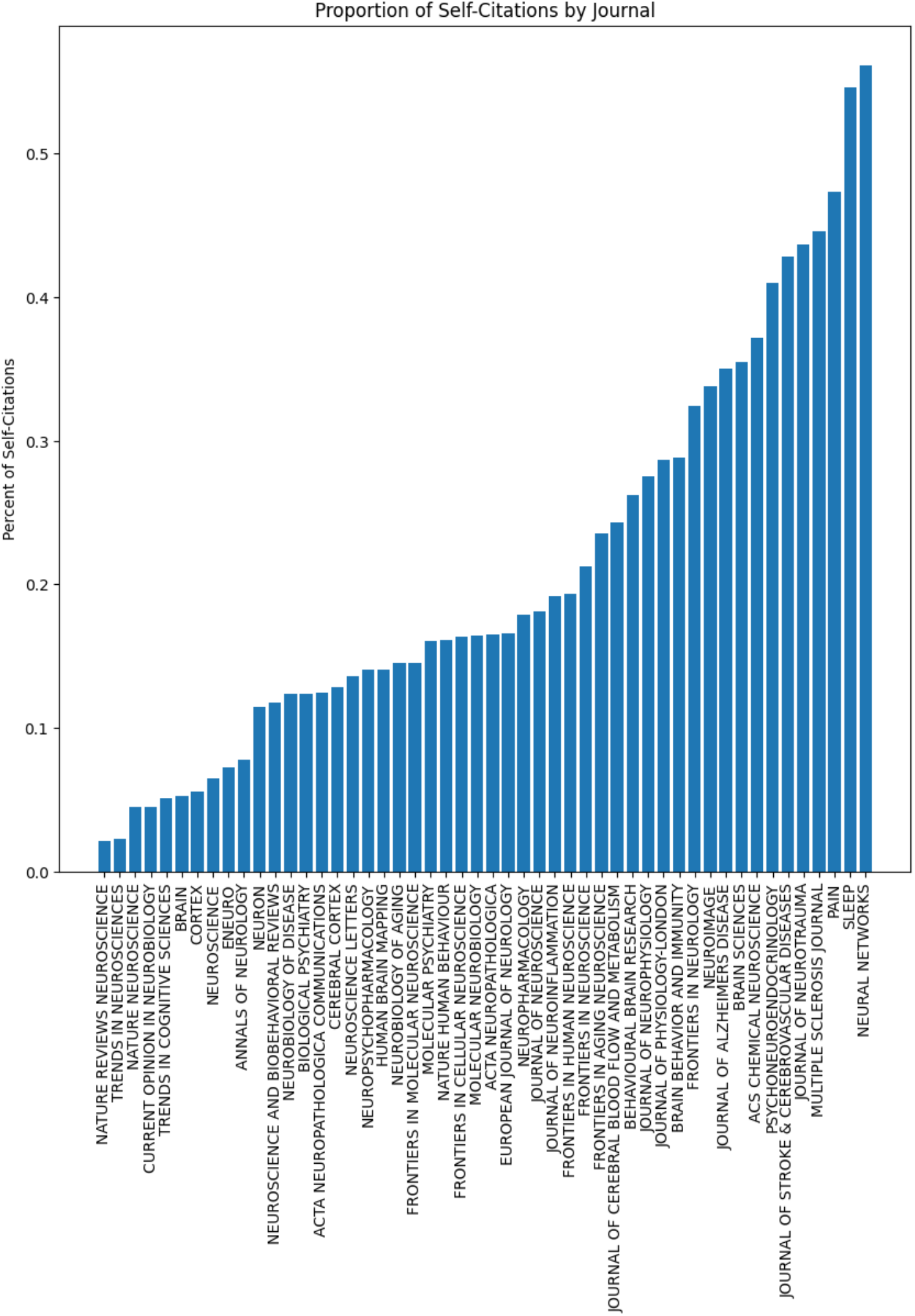
Percentage of citations made from a citing journal to the same journal.

### Full Data for Gender-based Breakdown

Here, we break down the gender-based citing behavior for each cluster.

**Extended Data Figure 7.**
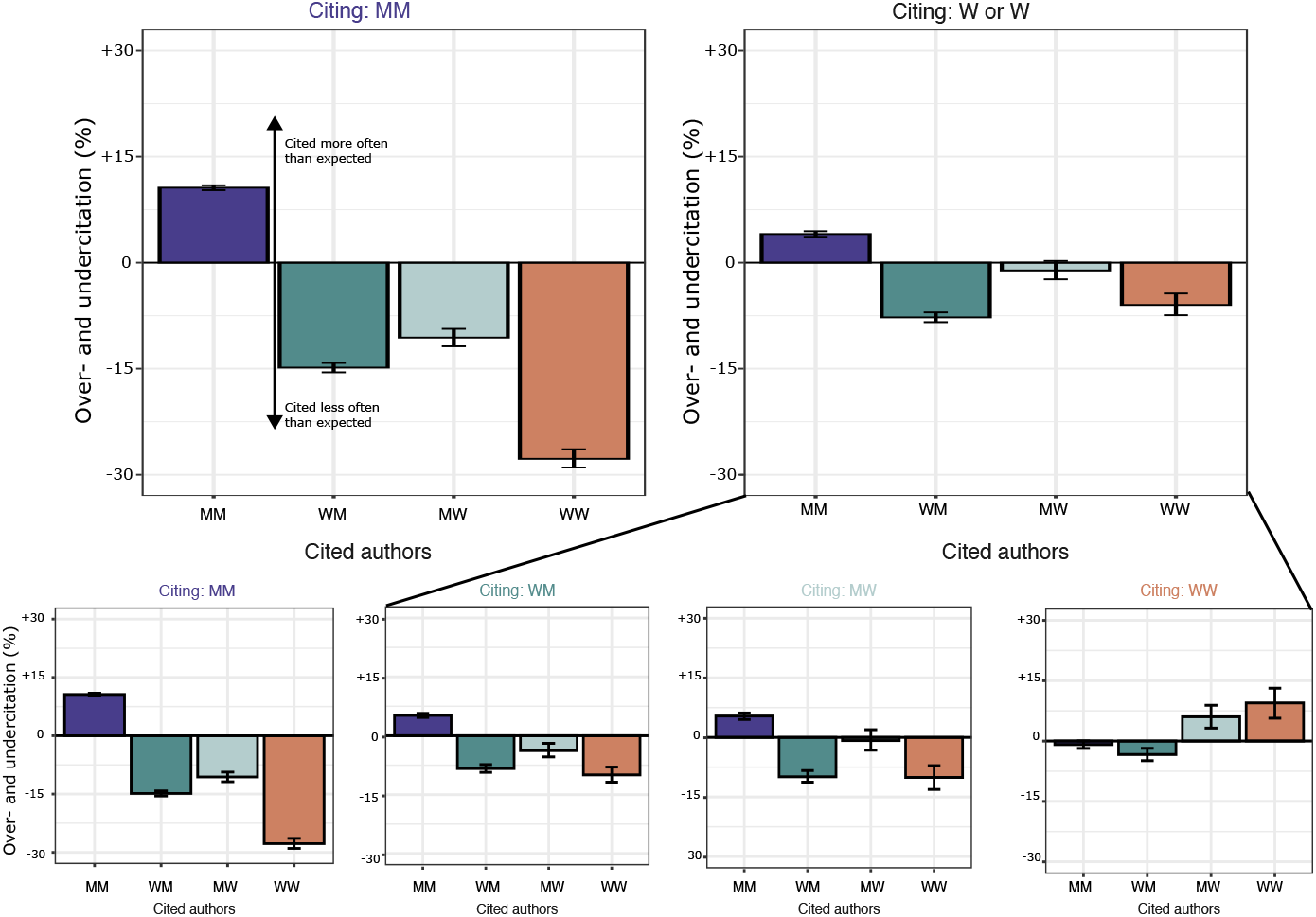
Effect of author gender on citation behavior within Group 1 papers. Degree of over- and undercitation of different author genders within MM and W∪W reference lists. **Top:** Papers with men as both first and last author overcite men to a greater extent than papers with women as either first or last author. **Bottom:** Full breakdown of gendered citation behavior within MM, WM, MW, and WW reference lists. Bars represent overall over/undercitation. Error bars represent the 95% CI of each over- and undercitation estimate, calculated from 500 bootstrap resampling iterations.

**Extended Data Figure 8.**
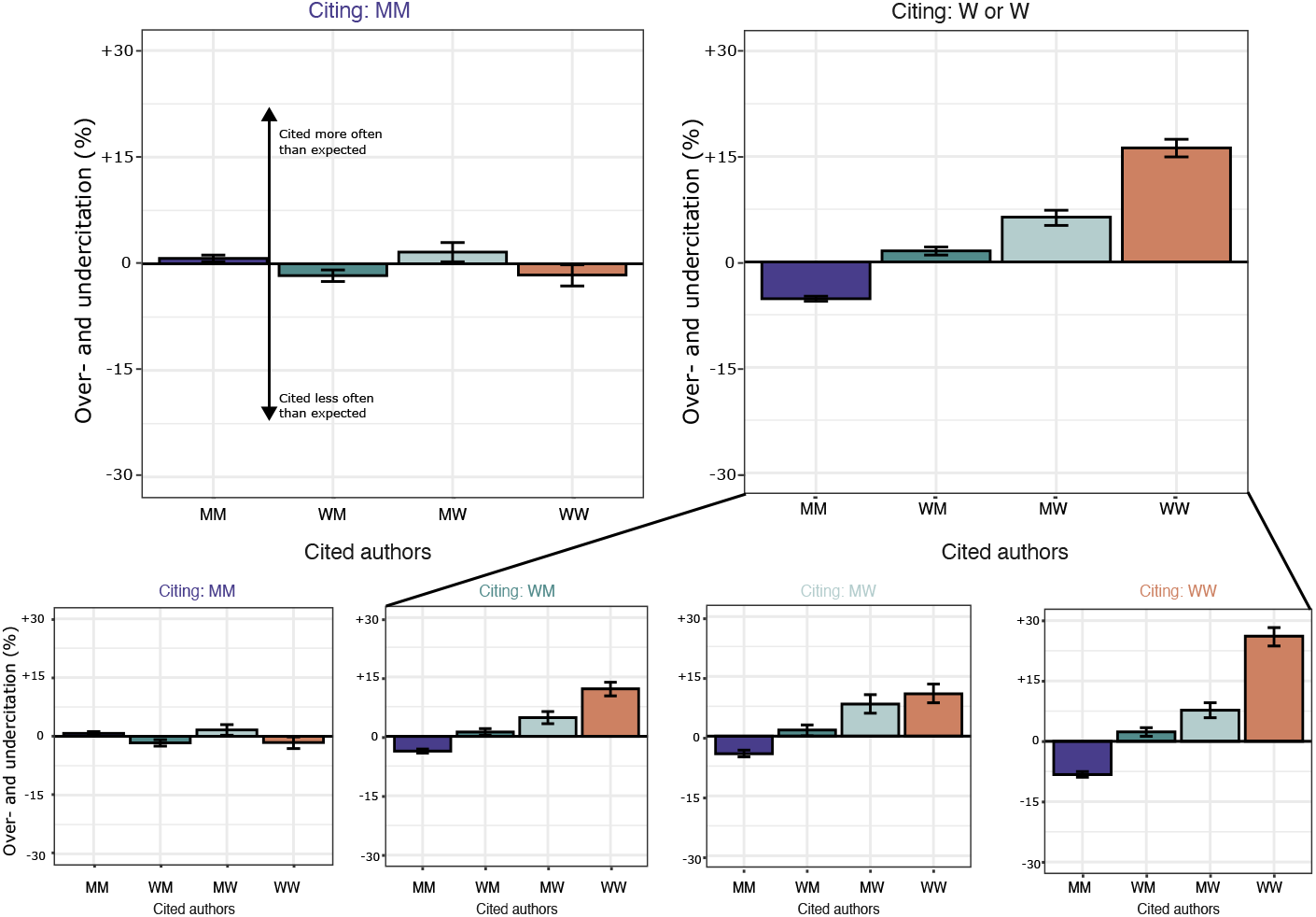
Effect of author gender on citation behavior within Group 2 papers. Degree of over- and undercitation of different author genders within MM and W∪W reference lists. **Top:** Papers with men as both first and last author cite in a way that is consistent with the proportion of genders in the field, while papers with at least one woman as an author overcite other papers with at least one woman author. **Bottom:** Full breakdown of gendered citation behavior within MM, WM, MW, and WW reference lists. Bars represent overall over/undercitation. Error bars represent the 95% CI of each over- and undercitation estimate, calculated from 500 bootstrap resampling iterations.

**Extended Data Figure 9.**
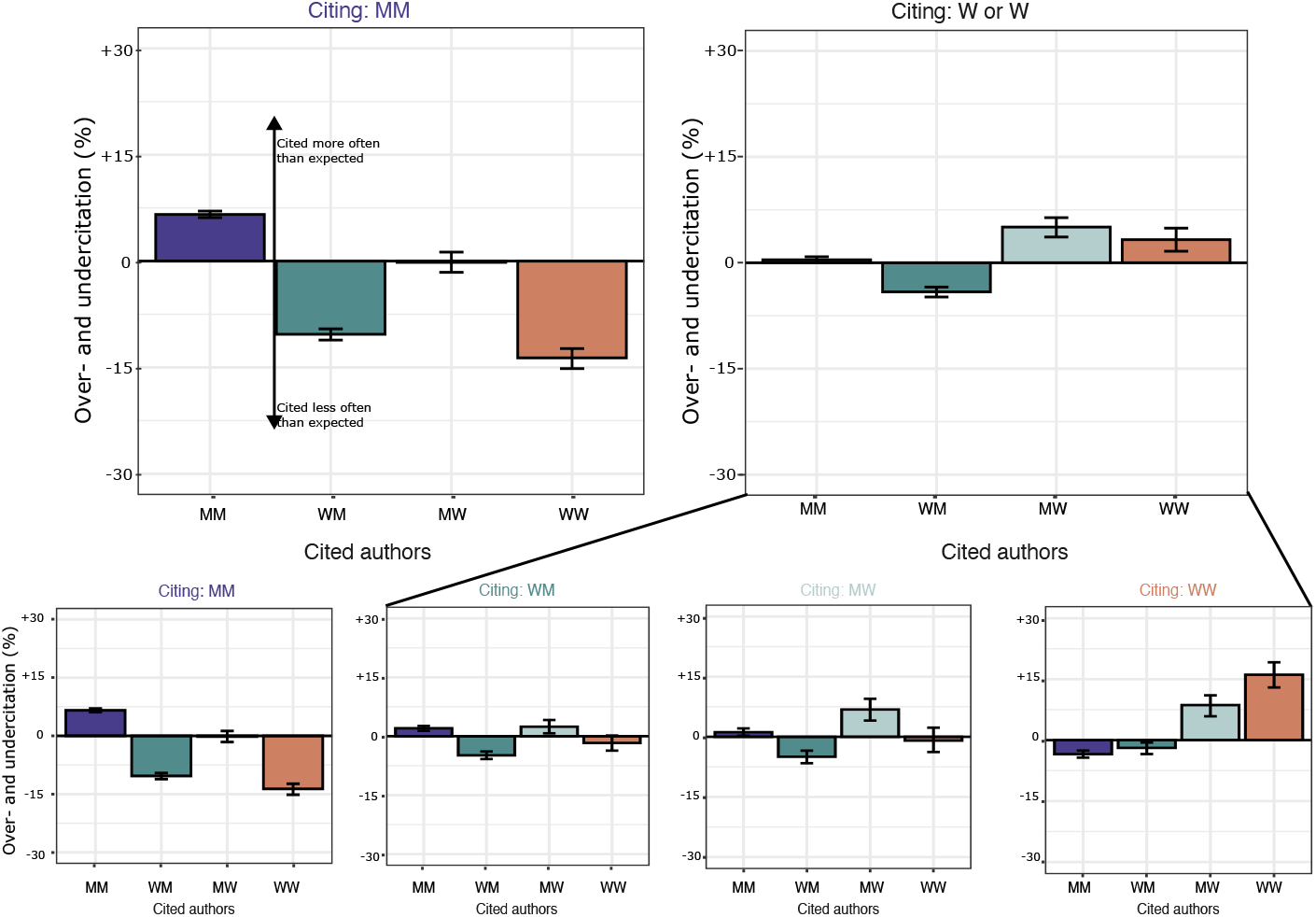
Effect of author gender on citation behavior within Group 3 papers. Degree of over- and undercitation of different author genders within MM and W∪W reference lists. **Top:** Papers with men as both first and last author overcite men to a greater extent than papers with women as either first or last author. **Bottom:** Full breakdown of gendered citation behavior within MM, WM, MW, and WW reference lists. Bars represent overall over/undercitation. Error bars represent the 95% CI of each over- and undercitation estimate, calculated from 500 bootstrap resampling iterations.

### Qualitative Interpretation of Group Topics

The role of neuroscience subfields is not directly accounted for within the primary analyses. Yet, we were curious whether our clustering method could reveal some relationship between subfield and citation behavior. We qualitatively explored this question by creating word clouds from the words in the titles of a subsample of every 100th article in each cluster using R’s *tm* (text-mining)(45) and *wordcloud2* (visualization) packages. Unsurprisingly, there was a great deal of overlap between clusters. However, there were also some qualitative differences: Group 1 seemed broadest, including both computational and microbiological methods, while Group 2 seemed more focused on wetlab disease-oriented neuroscience and microbiology, with an emphasis on rodent model organisms rather than humans. Finally, Group 3 appeared to be the most translational, focusing on humans and human-specific diseases. These tendencies could be investigated further with all papers (rather than every 100th), and could be complemented by quantitative analysis of topic frequency.

**Extended Data Figure 10.**
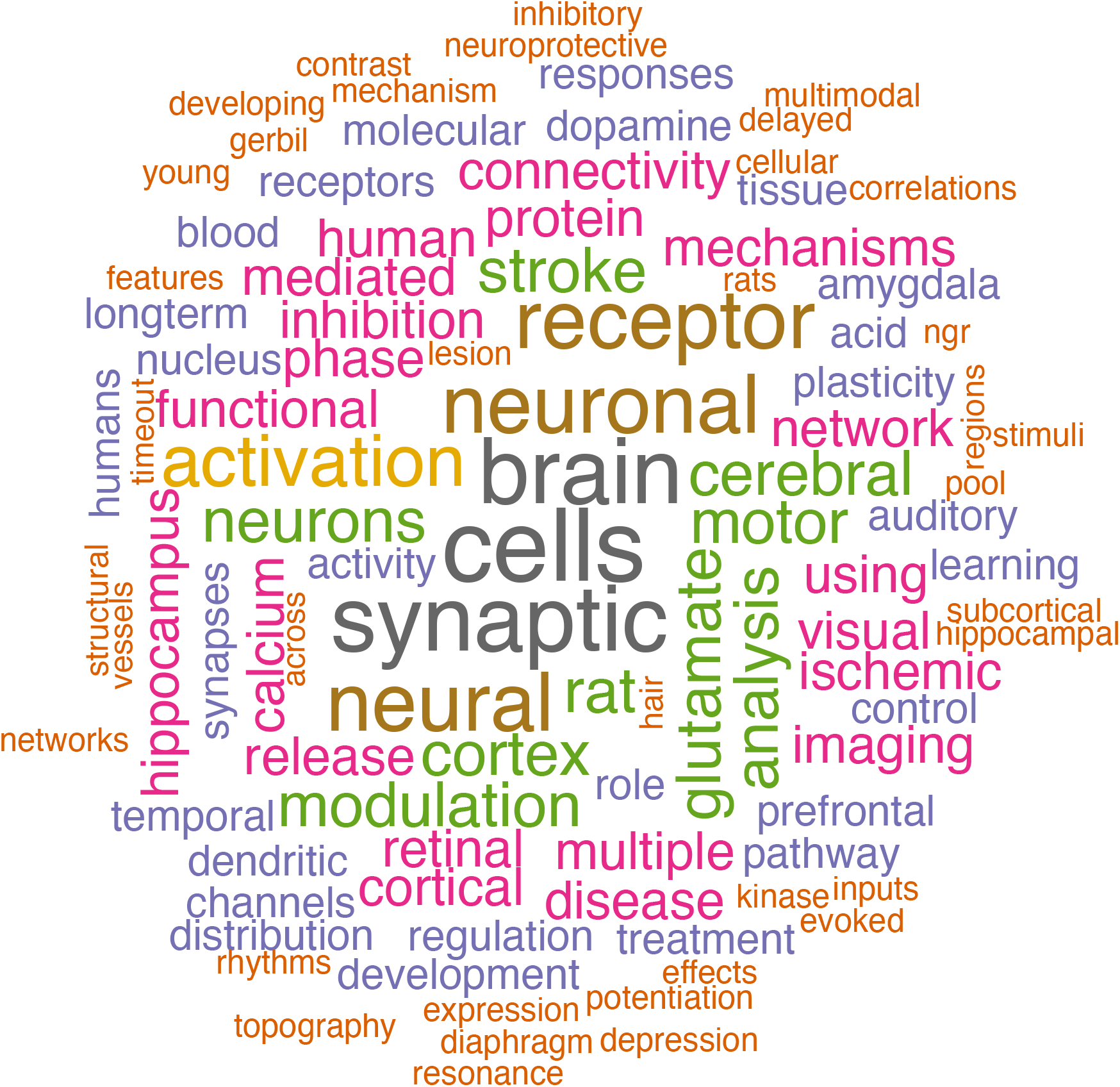
Word cloud created from a sample of the titles of Group 1 papers.

**Extended Data Figure 11.**
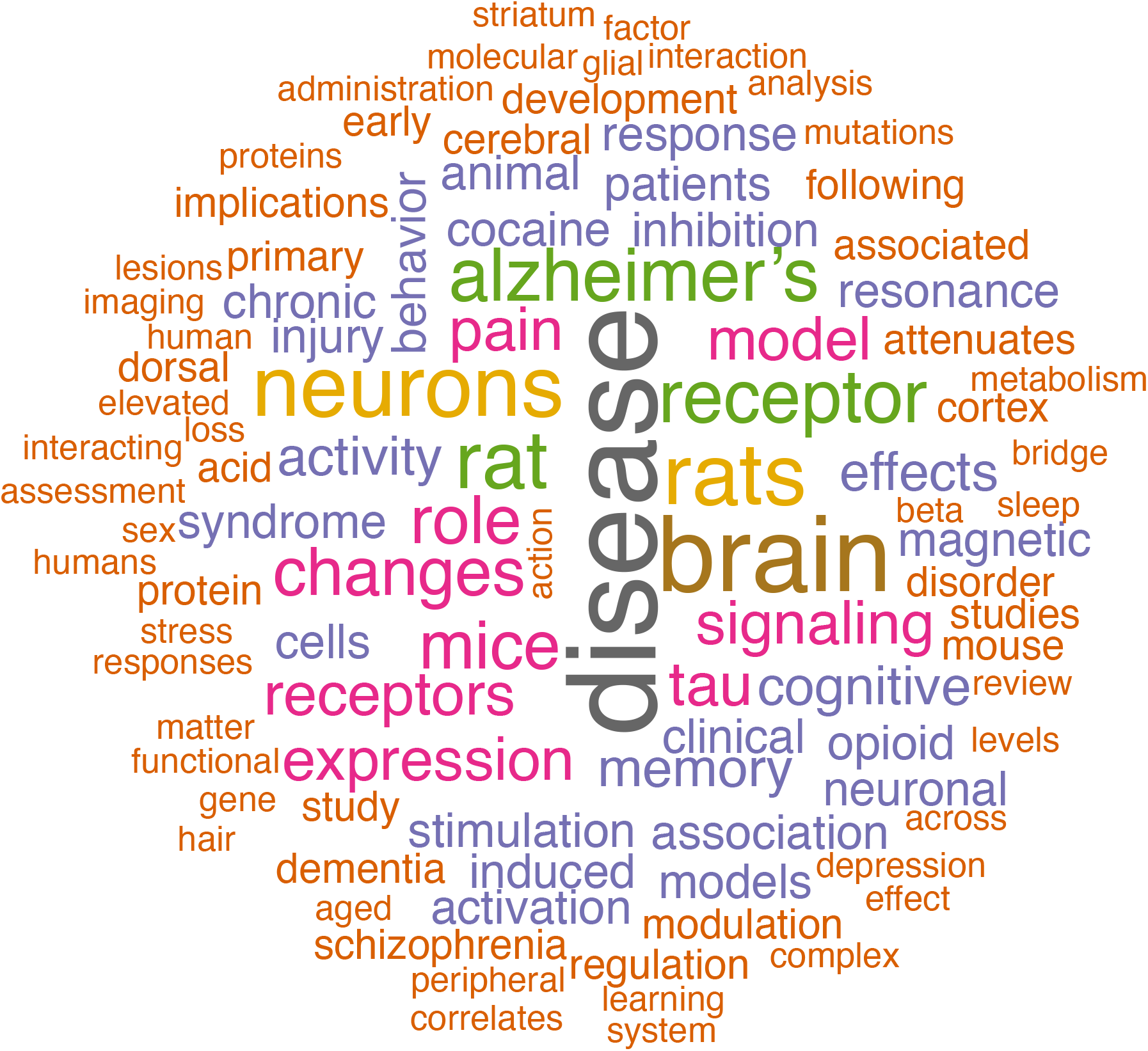
Word cloud created from a sample of the titles of Group 2 papers.

**Extended Data Figure 12.**
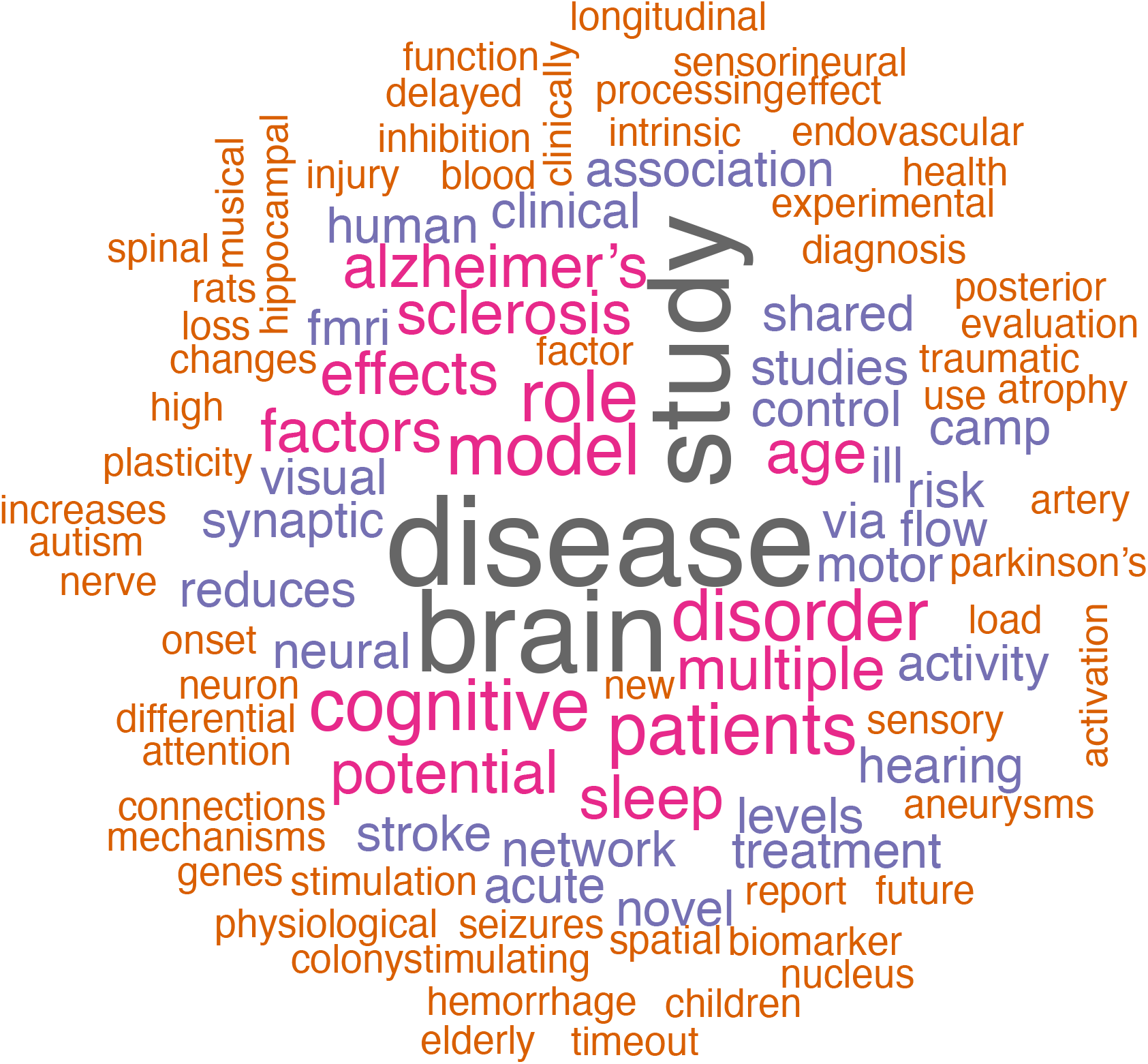
Word cloud created from a sample of the titles of Group 3 papers.

## Footnotes

1 Dworkin (13) did a sensitivity analysis on the effects of removing data for which both authors could not be specified and found that results did not significantly differ from assuming no gender imbalance in the missing entries.

2 Differences between the previous study and the current study are likely due to individual variation within journals: over the intervening period, *Brain* has been replaced by *Frontiers in Neuroscience* in the set of top five journals.

3 In our prior study, we conducted a sensitivity analyses by selecting a random sample of 200 authors, and found the accuracy of automated gender assignments to be 0.96% at the individual level, and 92% at the paper categorization level (13). In any event, errors in classification would bias results towards the null as they would break the link between gender and citation behavior. In addition, in our prior study we analyzed the effects of excluding data for which author gender was unknown. We imputed missing gender based on the GAM model, and did not observe a significant difference in citation gaps after imputation.

## Notes

https://github.com/jdwor/gendercitation

### Competing Interest Statement

The authors have declared no competing interest.

